# Inflammation-Induced Alternative Splicing in Human Endothelial Cells Reveals Genetic Mechanisms of Cardiovascular Disease Risk

**DOI:** 10.1101/2025.07.29.667484

**Authors:** Anna K. Golebiewski, Lindsey K. Stolze, Valentina D. Vazquez, Alhan Mehrabi Yazdi, Cecilia M. Careaga, Casey E. Romanoski

## Abstract

Alternative splicing modulates mRNA protein-coding sequence, stability, and translation rates, although it has not been comprehensively annotated in human endothelial cells (ECs). EC dysfunction is a hallmark of complex inflammatory diseases, including cancer and atherosclerosis. Therefore, this study modeled acute inflammation in vitro using 53 genetically distinct human aortic EC lines exposed to interleukin-1β (IL-1β) or control media. This approach identified 1,224 differentially spliced transcripts (DSTs) between IL-1β and control conditions. DSTs were enriched for alternative first (AF) exons, including several novel mRNA isoforms of disease-associated and metabolic genes. It was hypothesized and confirmed that AF splicing was driven by alternative promoters using ATAC-seq and ChIP-seq data. To identify alternative promoters driving IL-1β-dependent AF isoforms, a quantitative measure of promoter activity ratios was defined, and analysis found that histone 3 lysine 27 acetylation and binding of the transcription factors ERG and RELA often correlated with alternative promoter usage. Finally, the effect of common genetic variants on alternative first exon usage was interrogated through splicing quantitative trait locus (sQTL) analysis. Significant sQTLs were next submitted to genetic colocalization analysis with cardiovascular-related associations identified by genome-wide association studies (GWAS), finding colocalized signals at 66 human disease loci corresponding to 30 genes and 39 variants. These genetically regulated splicing differences provide plausible mechanisms explaining some of the genetic risk for cardiovascular-related diseases. Among the top signals are novel isoforms of Endothelial Protein C Receptor (PROCR) and Distal Membrane Arm Assembly Component 2 (DMAC2), whose splicing patterns colocalize with risk for coronary artery disease (CAD). This study demonstrates the prevalence of inducible alternative promoters and supports that ECs express numerous novel transcripts regulated by genetics and inflammation that are consistent with driving individual risk for cardiovascular disease.

**Graphical Abstract:** 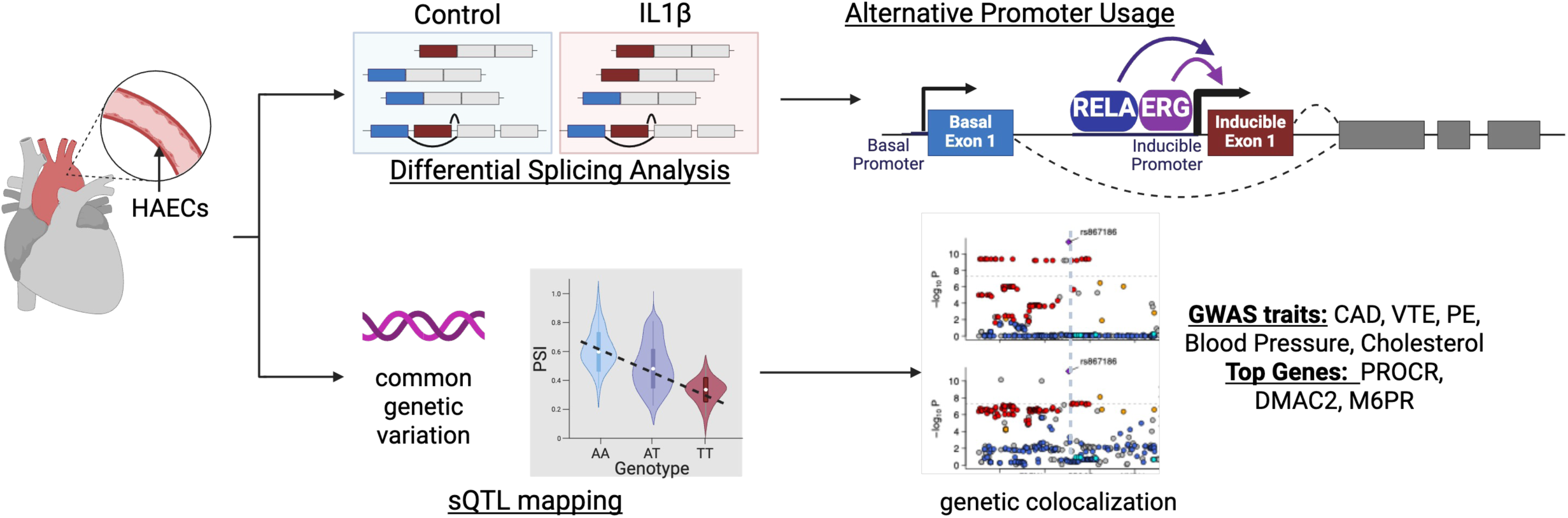

## Introduction

Endothelial cells (ECs) are found throughout the body in arteries, veins, and capillaries, where they play a central role in maintaining vascular tone and barrier function. Disruption of this barrier, referred to as endothelial dysfunction, is characterized by impaired nitric oxide synthesis, loss of barrier integrity, increased expression of inflammatory proteins, and enhanced recruitment of immune cells^1,2^. Since its initial characterization in 1986, endothelial dysfunction has been implicated in the pathogenesis of numerous complex diseases, including atherosclerosis, heart failure, diabetes, and kidney failure^1–3^. Pro-inflammatory stimuli such as tumor necrosis factor alpha (TNFα), lipopolysaccharide (LPS), and interleukin-1 beta (IL1β) induce an activated endothelial phenotype ^4^. Notably, IL1β blockade was shown to reduce recurrent heart attack rates in a clinical trial, highlighting the therapeutic potential of targeting inflammatory pathways^5^. To devise new strategies to combat endothelial dysfunction, it is essential to understand the diversity of molecular products generated by ECs in response to inflammatory stimulation such as IL1β.

**Alternative splicing (AS)** of RNAs is an important post-transcriptional process that allows a single gene to give rise to multiple mRNA and protein isoforms. More than 95% of multiexon human genes undergo AS, greatly expanding proteomic diversity ^6^. AS is orchestrated by the spliceosome complex that is recruited to pre-mRNAs by RNA-binding proteins (RBPs), transcription factors (TFs), and sequences in the RNA itself ^7^. AS results from different patterns of exon inclusion in final mRNAs through the removal of intronic sequences. AS can also arise from alternative transcriptional start and end sites. These processes can occur co-transcriptionally or post-transcriptionally, with co-transcriptional splicing generally producing more mature mRNA per pre-mRNA molecule ^8^.

Splicing outcomes are highly context- and cell type-specific ^9,10^, and their regulation is influenced by the spatial organization of the nucleus. Proximity to nuclear speckles, membraneless nuclear bodies enriched in splicing factors, enhances splicing efficiency and is governed by chromatin architecture, RNA polymerase II, and the coordinated action of RBPs and TFs ^11^. Notably, the induction signaling pathways, such as by transforming growth factor beta (TGFβ), have been shown to influence co-transcriptional splicing patterns ^8^, and inflammation-associated AS has been observed in macrophages and smooth muscle cells ^12,13^. However, transcriptome-wide characterization of AS in endothelial cells, particularly in response to inflammatory stimuli, remains limited. Recent studies demonstrate that *in vivo* both local inflammation from shear stress and immune-cells can induce AS in the endothelium^14,15^. Importantly, these AS genes are associated with important endothelial dysfunction-associated pathways including immune activation, cell junctions, and nuclear factor kabba B (NFkB) signaling^14^. This suggests that AS in endothelial cells plays a crucial role in endothelial dysfunction and likely cardiovascular disease and warrants further investigation.

Beyond AS, DNA polymorphisms that vary among people can affect phenotypes by affecting gene expression with cell type specificity. Quantitative Trait Locus (QTL) mapping, where genotypes across individuals in a population non-randomly associate with phenotypic differences, is a powerful method to identify genotype-phenotype relationships. At the molecular level, we and others have shown that QTL mapping of mRNA gene <u>e</u>xpression (termed eQTLs) are cell type specific. Specifically, we found that nearly 50% of EC eQTLs were specific to ECs and absent in the tissue-level eQTLS in GTEx (Genotype-Tissue Expression) database^16^. We hypothesize that a large proportion of alternative splicing elicited by IL1β and splicing QTLs (sQTLs), would also be specific to the EC cell type.

In this study, we sought to define the landscape of alternative splicing in ECs using an *in vitro* model of the acute response to the pro-inflammatory cytokine IL1β. Using transcriptomic and epigenomic data from 53 primary human aortic ECs (HAEC) lines, we identified differentially spliced genes (DSGs) upon IL1β treatment. We find that alternative promoter usage contributes to inflammation-induced AS and that the transcription factors NF-κB and ERG play complex roles in modulating transcript start sites from alternative promoters. We also utilized genetic variation in the EC cohort to identify sQTLs. We find that 87% of sQTLs were not identified by GTEx – the largest compendium of sQTLs available – thereby demonstrating that these sQTLS add to the existing breadth of transcriptomic diversity. Using genetic colocalization analysis, we identify 66 human disease loci corresponding to 30 AS genes (sGenes) and 39 genetic variants (sSNPs) where genetically driven splicing effects likely explain some of the genetic risk for cardiovascular-related diseases in genome-wide association studies (GWAS). Perhaps most interesting among these is the genetically regulated splicing pattern of *PROCR* (Endothelial protein C receptor; aka EPCR). At this locus, the established coronary artery disease (CAD)-risk allele rs867186-A, located in a 3’ exon of the gene, preferentially retains the rs867186-containing exon and untranslated region (UTR) in mRNA. In contrast, the CAD-protective rs867186-G allele more often excludes the canonical 3’ exon and instead retains a novel 3’ exon and UTR.

## RESULTS

### Alternative first exon splicing represents nearly one-third of AS in IL1β -treated HAECs

We quantified AS using RNA-seq data from 53 HAEC lines^16^ that were exposed *in vitro* to culture media containing IL1β (10 ng/mL) or control media for 4 hours (**Figure 1A**). Splicing effect size was assessed by Leafcutter^17^ using the change in percent spliced in (deltaPSI) metric that summarizes the difference in intron splicing between IL1β and control. At thresholds of 5% deltaPSI and 5% False Discovery Rate (FDR), we identified 1,224 Differentially Spliced Transcripts (DSTs) between control and IL1β (**Supp. Figure 1A, Supp. Table 1**). This corresponded to 288 Differentially Spliced Genes (DSGs), demonstrating that multiple transcript isoforms often arise from a single gene locus **(Supp. Figure 1A, Supp. Table 1).**

**Figure 1.**
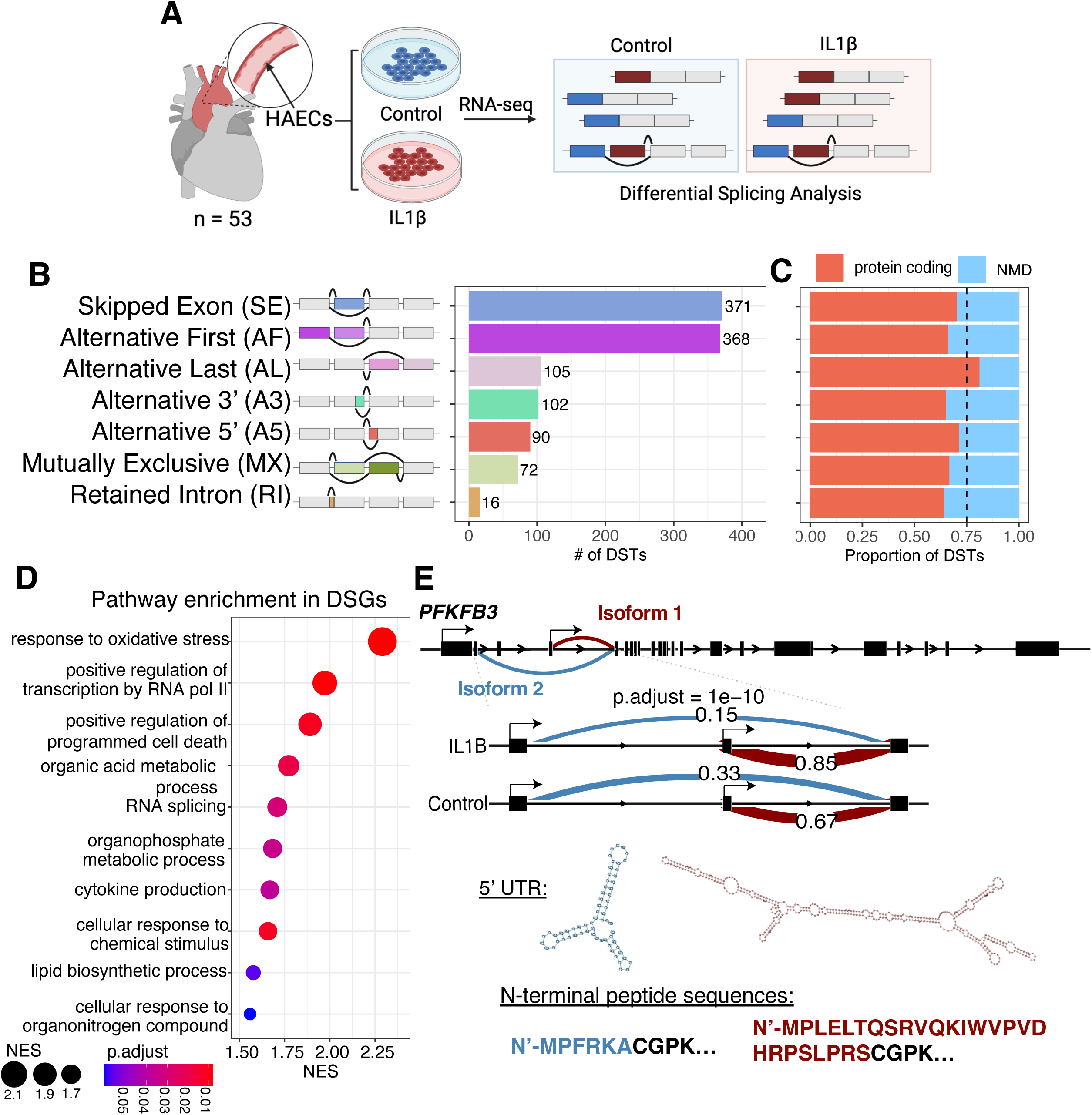
HAECs have a dynamic transcriptional response to IL1B via alternative splicing. A. 53 primary HAEC samples were treated with IL1B (10 ng/mL) and control media for four hours before collecting RNA. RNA-seq was performed, followed by differential splicing analysis using Leafcutter. B. Splice type categories in this study, and the number of significant DSTs (adjusted p.value < 0.05 and effect size (deltapsi) > 0.05, in each category. C. Percent of transcripts categorized as protein-coding or likely to be degraded by nonsense-mediated decay (NMD) based on RefSeq annotated stop codons. Dashed-line indicates average % of protein-coding transcripts (75%), including cryptic events (not shown). D. Pathway enrichment analysis was performed for DSGs, the top results are presented here. Full pathway enrichment analysis is available in Supplemental Table 2. E. PFKFB3 differential splicing at the first exon (AF). PFKFB3 has two transcription start sites (TSS’s) corresponding to mutually exclusive first exons for Isoform 1 (red) and Isoform 2 (blue). PSI values in IL1B and Control treatment are indicated on the sashimi plot for Isoform 1 and Isoform 2 first exon inclusion events. The PFKFB3 AF-transcripts have differing 5’ UTR (middle) and protein-coding sequences (bottom) in their respective first exons.

**Table 1:** sQTL and GWAS colocalization. Colocalizations between sQTLs and GWAS were assessed using coloc. SNPs were restricted to associations tested in both sQTL and GWAS studies. Colocalization tests were run for each AS intron separately to identify associations between AS and traits. Significance is defined as posterior probability of colocalization > 0.8. intron_PP.H4.abf—posterior probability of colocalization at a given intron.

| Gene | SNP | Intron | GWAS |  | Control sQTL |  | IL1B sQTL |  | Probability |
| --- | --- | --- | --- | --- | --- | --- | --- | --- | --- |
|  |  |  | gwas_pvalue | gwas_beta | FDR_Control | Beta_Control | FDR_IL1B | Beta_IL1B | intron_PP.H4.abf |
| Coronary Artery Disease |  |  |  |  |  |  |  |  |  |
| PROCR | rs867186 | 20:35176446:35215893:clu_14879_NA | 6.84e-12 | -0.05730 | 2.65e-14 | 0.0649037 | 1.9e-13 | 0.0598204 | 0.9999962 |
| PROCR | rs867186 | 20:35176446:35176698:clu_14879_NA | 6.84e-12 | -0.05730 | 1.14e-14 | -0.0766561 | 2.09e-15 | -0.0716977 | 1.0000000 |
| TMEM91 | rs7259900 | 19:41382921:41383886:clu_8457_NA | 6.29e-10 | -0.03560 | 0.000127 | 0.1625228 | NA | NA | 0.8323292 |
| ATP5SL | rs1043413 | 19:41432408:41433537:clu_8461_NA | 3.66e-08 | 0.03150 | 6e-11 | -0.1043871 | 4.38e-09 | -0.0859710 | 0.9941839 |
| HLA-B | rs1140546 | 6:31354665:31355107:clu_13338_NA | 1.51e-06 | 0.03340 | 0.00348 | 0.0753015 | 0.0285 | 0.0649257 | 0.8948707 |
| PARP12 | rs2286196 | 7:140026348:140028613:clu_16311_NA | 8.29e-06 | -0.03160 | 0.000153 | 0.0911376 | 2.77e-11 | 0.1044265 | 0.9559289 |
| ERAP2 | rs55770741 | 5:96880859:96883792:clu_7392_NA | 1.38e-05 | -0.02520 | 1.06e-05 | -0.0468416 | 1.9e-05 | -0.0281599 | 0.8158660 |
| APOPT1 | rs71417868 | 14:103563124:103587274:clu_3906_NA | 2.71e-05 | 0.02650 | 0.000532 | -0.0324354 | 6.95e-06 | -0.0481726 | 0.8757482 |
| Deep Vein Thrombosis |  |  |  |  |  |  |  |  |  |
| PROCR | rs867186 | 20:35176446:35176698:clu_14879_NA | 4.48e-09 | 0.15980 | 1.14e-14 | -0.0766561 | 2.09e-15 | -0.0716977 | 0.9606076 |
| PROCR | rs867186 | 20:35176446:35215893:clu_14879_NA | 4.48e-09 | 0.15980 | 2.65e-14 | 0.0649037 | 1.9e-13 | 0.0598204 | 0.9606075 |
| ARL6IP4 | rs4275659 | 12:122981299:122981571:clu_11263_NA | 2.72e-05 | 0.07243 | 4.06e-07 | 0.0463675 | 4.44e-05 | 0.0444106 | 0.9577743 |
| PSORS1C1 | rs562775931 | 6:31114891:31125739:clu_13339_NA | 2.72e-05 | -0.07181 | 0.0382 | -0.2374057 | 6.18e-06 | -0.3180288 | 0.9157402 |
| PSORS1C1 | rs562775931 | 6:31114891:31115136:clu_13339_NA | 2.72e-05 | -0.07181 | 0.0447 | 0.1925162 | 6.67e-05 | 0.2553488 | 0.9151033 |
| Diastolic Blood Pressure |  |  |  |  |  |  |  |  |  |
| ERAP2 | rs1559359 | 5:96900245:96901506:clu_7394_NA | 1.58e-11 | -0.01661 | 9.49e-06 | -0.2209348 | 0.00236 | -0.1936838 | 0.9210396 |
| ERAP2 | rs1559359 | 5:96880859:96883792:clu_7392_NA | 1.58e-11 | -0.01661 | 1.06e-05 | -0.0475170 | 2.51e-05 | -0.0288766 | 0.8991886 |
| LTBP4 | rs1051481 | 19:40617020:40617100:clu_8436_NA | 7.55e-10 | 0.02000 | 0.00328 | 0.0729420 | NA | NA | 0.8130061 |
| GRB10 | rs10226465 | 7:50727890:50755887:clu_15910_NA | 1.82e-09 | -0.01464 | NA | NA | 0.00535 | 0.0068215 | 0.8504497 |
| RBM23 | rs28600251 | 14:22905453:22906195:clu_3457_NA | 3.76e-07 | 0.01262 | 0.000123 | -0.0660288 | 0.00131 | -0.0544379 | 0.8613282 |
| LPP | rs13076750 | 3:188341719:188406112:clu_7032_NA | 4.46e-07 | 0.01407 | 3.22e-07 | 0.2403225 | 2.16e-13 | 0.3173935 | 0.9953968 |
| LPP | rs13076750 | 3:188225527:188406112:clu_7032_NA | 4.46e-07 | 0.01407 | 1.11e-06 | -0.2678076 | 1.29e-07 | -0.2871473 | 0.9953828 |
| LPP | rs13076750 | 3:188225527:188341656:clu_7032_NA | 4.46e-07 | 0.01407 | 0.00175 | 0.0677887 | 0.00148 | 0.0533583 | 0.9644933 |
| RBM23 | rs61977741 | 14:22905453:22906195:clu_3457_NA | 2.68e-06 | -0.01156 | 2.61e-05 | 0.0844795 | 9.4e-07 | 0.0860142 | 0.8933048 |
| M6PR | rs149871778 | 12:8946387:8949488:clu_10519_NA | 6.38e-06 | 0.01792 | 3e-04 | 0.1038984 | 2.13e-06 | 0.1461048 | 0.9715790 |
| M6PR | rs149871778 | 12:8946405:8949488:clu_10519_NA | 6.38e-06 | 0.01792 | 0.000254 | -0.1037522 | 3.01e-06 | -0.1419965 | 0.9676309 |
| M6PR | rs4883201 | 12:8946387:8949488:clu_10519_NA | 2.3e-05 | 0.01682 | 2.84e-07 | 0.1099606 | 5.72e-06 | 0.1400069 | 0.9393221 |
| M6PR | rs4883201 | 12:8946405:8949488:clu_10519_NA | 2.3e-05 | 0.01682 | 2.76e-07 | -0.1092253 | 8.34e-06 | -0.1358751 | 0.9393228 |
| HDL-Cholesterol |  |  |  |  |  |  |  |  |  |
| ARL6IP4 | rs4275659 | 12:122981299:122981571:clu_11263_NA | 8.51e-33 | -0.03363 | 4.06e-07 | 0.0463675 | 4.44e-05 | 0.0444106 | 0.9930257 |
| M6PR | rs149871778 | 12:8946387:8949488:clu_10519_NA | 6.31e-18 | -0.03564 | 3e-04 | 0.1038984 | 2.13e-06 | 0.1461048 | 0.9333041 |
| M6PR | rs149871778 | 12:8946405:8949488:clu_10519_NA | 6.31e-18 | -0.03564 | 0.000254 | -0.1037522 | 3.01e-06 | -0.1419965 | 0.9241181 |
| OASL | rs7979478 | 12:121027817:121031442:clu_11221_NA | 8.32e-16 | -0.02112 | 0.00116 | -0.1902392 | NA | NA | 0.8326739 |
| RSPRY1 | rs141312419 | 16:57186785:57204504:clu_9338_NA | 3.24e-15 | 0.02012 | NA | NA | 5.27e-05 | 0.0745889 | 0.9783969 |
| RSPRY1 | rs141312419 | 16:57186451:57204504:clu_9338_NA | 3.24e-15 | 0.02012 | NA | NA | 0.00191 | -0.0739219 | 0.8873397 |
| SH3YL1 | rs17714252 | 2:219001:229966:clu_1_NA | 4.17e-14 | 0.02015 | 8.13e-06 | 0.2083735 | 0.0294 | 0.0982292 | 0.8643034 |
| HYAL3 | rs2282749 | 3:50297672:50299213:clu_6471_NA | 1.05e-13 | -0.01905 | NA | NA | 0.00601 | -0.0992046 | 0.8115889 |
| THOC5 | rs2074948 | 22:29529239:29531831:clu_4119_NA | 7.76e-12 | -0.02062 | 0.00123 | 0.1345806 | 7.54e-05 | 0.1494768 | 0.8974384 |
| RP11-465B22.3 | rs9442388 | 1:1063201:1065830:clu_1098_NA | 2.17e-06 | -0.01220 | NA | NA | 0.00018 | -0.3208501 | 0.8013532 |
| CPNE1 | rs2425084 | 20:35626381:35626735:clu_14886_NA | 5.02e-06 | -0.01718 | 0.00512 | 0.0439686 | 1.69e-05 | 0.0402806 | 0.8542489 |
| LDL-Cholesterol |  |  |  |  |  |  |  |  |  |
| ERCC1 | rs7247937 | 19:45409725:45416821:clu_8550_NA | 6.61e-21 | -0.02643 | NA | NA | 0.0325 | 0.0074686 | 0.8425422 |
| TSPAN4 | rs4963153 | 11:843050:847201:clu_4480_NA | 2.05e-08 | -0.01425 | NA | NA | 0.000829 | -0.0402756 | 0.8277492 |
| RMDN1 | rs34499209 | 8:86472479:86474820:clu_15435_NA | 2.56e-08 | 0.01731 | 1.93e-11 | -0.2049788 | 1.02e-07 | -0.1649761 | 0.9733340 |
| RMDN1 | rs56810922 | 8:86474358:86474820:clu_15435_NA | 1.1e-07 | -0.01555 | 2.75e-11 | -0.2206454 | 2.07e-10 | -0.2156722 | 0.8601691 |
| RMDN1 | rs56810922 | 8:86472479:86474820:clu_15435_NA | 1.1e-07 | -0.01555 | 1.93e-11 | 0.2100304 | 5.09e-09 | 0.1853978 | 0.8512618 |
| RMDN1 | rs373220036 | 8:86480332:86486484:clu_15437_NA | 1.26e-07 | 0.01543 | NA | NA | 2.53e-05 | 0.0819766 | 0.8466691 |
| RER1 | rs6671730 | 1:2394728:2395784:clu_1149_NA | 2.28e-07 | -0.01317 | NA | NA | 0.00844 | 0.0152291 | 0.8060775 |
| RAB8B | rs61135268 | 15:63189748:63223848:clu_9892_NA | 5.42e-07 | 0.01467 | NA | NA | 4.33e-05 | 0.0642284 | 0.9916646 |
| NOD1 | rs2736723 | 7:30463698:30478606:clu_15821_NA | 1.02e-06 | -0.01247 | 0.029 | -0.1649409 | 0.000447 | -0.2116788 | 0.9331191 |
| STXBP4 | rs2541236 | 17:54999451:54999626:clu_12820_NA | 3.69e-06 | -0.01283 | 1.18e-10 | -0.3516956 | 2.32e-10 | -0.3214674 | 0.9520016 |
| STXBP4 | rs2541236 | 17:54999451:54999632:clu_12820_NA | 3.69e-06 | -0.01283 | 2.5e-10 | 0.2992596 | 1.22e-08 | 0.2554486 | 0.9520015 |
| NOD1 | rs2709801 | 7:30460041:30478606:clu_15821_NA | 4.29e-06 | -0.01171 | 0.000217 | 0.3031498 | 0.000137 | 0.3042899 | 0.9367356 |
| NOD1 | rs1558067 | 7:30460041:30478606:clu_15821_NA | 6.87e-06 | -0.01141 | 0.000287 | 0.2469521 | 5.31e-05 | 0.2715466 | 0.9561251 |
| Pulmonary Embolism |  |  |  |  |  |  |  |  |  |
| PSORS1C1 | rs562470954 | 6:31114891:31125739:clu_13339_NA | 0.000422 | -0.09385 | 0.0382 | -0.2374057 | 6.18e-06 | -0.3180288 | 0.8330205 |
| PSORS1C1 | rs562470954 | 6:31114891:31115136:clu_13339_NA | 0.000422 | -0.09385 | 0.0447 | 0.1925162 | 6.67e-05 | 0.2553488 | 0.8362819 |
| Systolic Blood Pressure |  |  |  |  |  |  |  |  |  |
| BAG6 | rs396369 | 6:31651776:31652424:clu_13364_NA | 9.55e-25 | 0.02678 | NA | NA | 0.0164 | 0.0651512 | 0.8426626 |
| BAG6 | rs1052486 | 6:31648751:31648911:clu_13363_NA | 3.47e-24 | 0.02343 | 0.00055 | -0.0485878 | NA | NA | 0.9050450 |
| BAG6 | rs1052486 | 6:31648964:31649199:clu_13363_NA | 3.47e-24 | 0.02343 | 0.00158 | 0.0469653 | NA | NA | 0.8102605 |
| RBM23 | rs28600251 | 14:22905453:22906195:clu_3457_NA | 5.46e-09 | 0.01376 | 0.000123 | -0.0660288 | 0.00131 | -0.0544379 | 0.8472132 |
| RBM23 | rs61977741 | 14:22905453:22906195:clu_3457_NA | 2.4e-08 | -0.01305 | 2.61e-05 | 0.0844795 | 9.4e-07 | 0.0860142 | 0.8825732 |
| M6PR | rs149871778 | 12:8946387:8949488:clu_10519_NA | 5.79e-05 | 0.01508 | 3e-04 | 0.1038984 | 2.13e-06 | 0.1461048 | 0.8234604 |
| M6PR | rs149871778 | 12:8946405:8949488:clu_10519_NA | 5.79e-05 | 0.01508 | 0.000254 | -0.1037522 | 3.01e-06 | -0.1419965 | 0.8028618 |
| Total Cholesterol |  |  |  |  |  |  |  |  |  |
| M6PR | rs149871778 | 12:8946387:8949488:clu_10519_NA | 4.47e-15 | -0.03080 | 3e-04 | 0.1038984 | 2.13e-06 | 0.1461048 | 0.9579449 |
| M6PR | rs149871778 | 12:8946405:8949488:clu_10519_NA | 4.47e-15 | -0.03080 | 0.000254 | -0.1037522 | 3.01e-06 | -0.1419965 | 0.9520184 |
| M6PR | rs4883201 | 12:8946387:8949488:clu_10519_NA | 1.86e-14 | -0.03011 | 2.84e-07 | 0.1099606 | 5.72e-06 | 0.1400069 | 0.8996619 |
| M6PR | rs4883201 | 12:8946405:8949488:clu_10519_NA | 1.86e-14 | -0.03011 | 2.76e-07 | -0.1092253 | 8.34e-06 | -0.1358751 | 0.8996632 |
| ERCC1 | rs7247937 | 19:45409725:45416821:clu_8550_NA | 2.04e-14 | -0.02057 | NA | NA | 0.0325 | 0.0074686 | 0.8661606 |
| RMDN1 | rs34499209 | 8:86472479:86474820:clu_15435_NA | 5.82e-06 | 0.01346 | 1.93e-11 | -0.2049788 | 1.02e-07 | -0.1649761 | 0.9607242 |
| UAP1 | rs6672561 | 1:162590511:162597792:clu_2327_NA | 4.51e-05 | -0.01170 | 5.19e-06 | -0.1748764 | 0.000381 | -0.1558409 | 0.8517596 |

We partitioned splice junctions into the following categories: Skipped Exons (SE), Retained Introns (RI), Mutually eXclusive (MX) exons, Alternate First (AF) exons, Alternative Last (AL) exons, Alternative 3’ splice sites (A3), and Alternative 5’ splice sites (A5) (**Figure 1B, Supp. Figure 1B**). Interestingly, we found 120 cryptic junctions whose annotations were missing from the reference transcriptome. All junctions identified are in **Supplemental Table 1**. 75% of the DSTs are predicted to effect protein-coding exons and 25% are predicted to elicit nonsense-mediated decay (NMD) (**Figure 1B**), suggesting for the most part that IL1β DSTs are likely to impact protein structure and or abundance.

Among splice types, AF splices had the largest effect sizes relative to the others (**Supp. Figure 1C**) and were enriched in the significant list of DSTs (p = 1 × 10^−25^) relative to the expected frequency of AF splices annotated in the transcriptome (**Figure 1C**). A3 splice events were also enriched, but at lower frequency and average effect size. We compared the expected proportion of transcript types assessed for differential gene expression and found that differentially expressed genes (DEGs) are significantly enriched for genes that are also AF-DSGs or SE-DSGs (p = 7.14e^−9^, p = 3.4e^−4^). However, both AF and SE DSTs have relatively modest effect sizes for differential expression, indicating that the differential splicing and expression responses may be regulated by separate mechanisms (**Supp. Figure 1C**). A similar study in monocyte-derived macrophages found that AF splicing was enriched in response to inflammatory stimulation^18^. Comparing the EC and macrophage ^18^ responses, we found that greater than 95% of DSTs are unique to either ECs or macrophages demonstrating the specificity of responses (**Supp Figure 1E**).

### Pathway enrichment of DSGs identifies metabolic and inflammatory pathways

Overall, DSGs were statistically enriched for inflammation associated pathways, including ‘Response to Oxidative Stress’, ‘Positive Regulation of Programmed Cell Death’, ‘Cytokine Production’, and ‘Cellular Response to Chemical Stimulus’ (**Figure 1D, Supp. Table 2**). Metabolic processes were also enriched, as reflected by the terms, ‘Organic Acid Metabolic Process’, ‘Organophosphate Metabolic Process’, ‘Lipid Biosynthetic Process’, and ‘Cellular Response to Organonitrogen Compound’ (**Figure 1D, Supp. Table 2**). Together, these findings show that AS is a prominent mechanism governing the HAECs gene expression response to IL1β. Furthermore, AF-DSGs are overrepresented in gene sets of important biological pathways insofar as AF-DSGs make up only 23% of all DSGs but comprise 60% of genes driving pathway enrichment (**Supp. Figure 1F**). For example, AF-DSGs represent 6 of the 7 DSGs in the oxidative stress pathway (*ABL1, GPX4, KDM6B, NCOA7, RCAN1, and SESN1*) (**Supp. Table 2**).

### Differential splicing analysis identifies novel AS of *PFKFB3*

An interesting example of IL1β-responsive AS is 6-phosphofructo-2-kinase/fructose-2,6-biphosphatase 3 (PFKFB3). *PFKFB3* is an AF-DSG that is both differentially spliced and differentially expressed with IL1β (deltaPSI = 0.245, p.adj = 1 × 10^−10^; log_2_FC = 1.057, FDR = 1.29 × 10^−31^) (**Figure 1E**). PFKFB3 is a rate-limiting enzyme in the glycolysis metabolic pathway. It phosphorylates fructose-6-phosphate to produce fructose-2,6-bisphosphate, an allosteric activator of PhosphoFructoKinase-1 (PFK1)^19,20^. AS of *PFKFB3* produces the canonical *PFKFB3* (Isoform 1) which is upregulated by IL1β treatment, and a novel transcript (Isoform 2) that is less abundant than Isoform 1. In control conditions, 33% of *PFKFB3* mRNAs contain the Isoform 2-specific splice, and after IL1β exposure, its proportion decreases to 15% (**Figure 1E**). The two isoforms differ only by their first exon, which results in a change in the 5’ UTR and the N-terminal protein sequence (**Figure 1E**).

We employed quantitative real-time PCR with isoform-specific primers to evaluate relative expression levels of *PFKFB3* isoforms upon 4, 8, and 24 hours of IL1β treatment in HAECs. Consistent with the RNA-seq at 4 hours, both isoforms were significantly increased in expression after 4 hours by different magnitudes. *PFKFB3* isoform 1 RNA increased six**-**fold (p = 0.0019) whereas *PFKFB3* isoform 2 increased by less than two-fold (p = 0.023) (**Supp Figure 1G**). After 8 hours, PFKFB3 isoform 2 was no longer upregulated (p = 0.11), while *PFKFB3* isoform 1 remained elevated (p = 0.0078) until 24 hours when levels came down (isoform 1: p = 0.064; isoform 2: p = 0.029) (**Supp Figure 1G**). Beyond being an important metabolic enzyme, the pattern of AS at the *PFKFB3* locus represents an interesting use of first exons and alternative promoters that is initiated by inflammatory conditions and may be applicable to other AF-DSGs.

### Defining alternative promoters at AF transcripts

We sought to delve deeper into how AF-DSGs are regulated, and whether alternative promoters drive their transcription. Promoters are dynamic regions of DNA upstream of gene transcription start sites (TSS) where transcription factors (TFs) recruit RNA polymerase to initiate transcription. We distinguished alternative promoters in this study into two categories: ***inducible promoters (P_inducible_)*** and ***basal promoters (P_basal_).*** P_inducible_ are regions directly 5’ to alternative first exons containing more RNA-seq reads in IL1β treatment relative to RNAs measured in control-treated HAECs. In other words, IL1β treatment ‘induces’ promoter activity and transcription across the alternative first exon. In contrast, P_basal_ defines promoters immediately 5’ to alternative first exons that express more RNA in control (basal) conditions relative to IL1β treatment (**Figure 2A**). Only DSGs with paired AF-DSTs were considered to have P_inducible_ and P_basal,_ more complex systems with more than two AF exons or only one significant AF exon were not considered in the following analyses.

**Figure 2.**
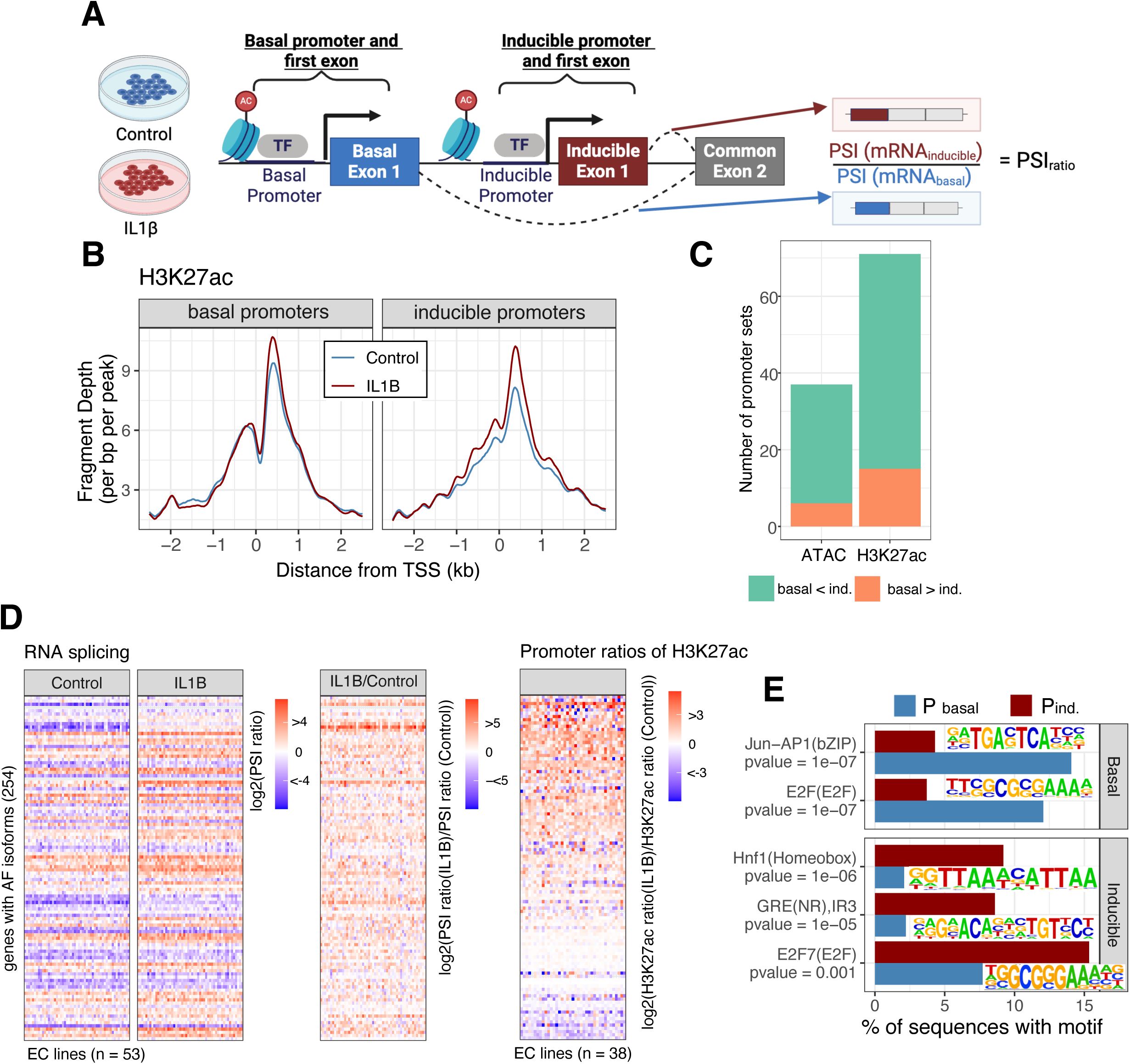
Alternative Promoter Usage Drives Alternative First Exon Expression. A. We defined basal first exons and promoters (P_basal_) as having increased PSI for adjoining intron in control treatment, and inducible first exons and promoters (P_inducible_) as having increased PSI for adjoining introns in IL1B treatment. PSI_ratio_ is then defined as the mRNA_inducible_ to the mRNA_basal_. B. ChIP-seq for H3K27ac at P_basal_ and P_inducible_ for Control (blue) and IL1B (red) treated HAECs (n = 50). Peaks are centered on the transcription start site (TSS) and distance is plotted as the distance from the TSS in kilobases (kb). C. The PSI_ratio_ in Control and IL1B treatments, and the ratio of the two PSI_ratio_ values to show the effect of treatment. By definition, all log2(PSI_ratio_) values are > 0 because there is more mRNA_inducible_ than mRNA_basal_. This is compared to the ratio of the two PARs for H3K27ac (= tag count in promoter region) at P_inducible_ to P_basal_ for Control and IL1B treatments to show the effect of treatment on the PAR. D. Number of significant differences between PAR_IL1B_ and PAR_Control_ for H3K27ac and ATAC. E. Motif enrichment in P_basal_ (top) and P_inducible_ (bottom) for known motif sets in the Homer database.

### H3K27ac is a good predictor of IL1β -driven alternative promoters

Next, to interrogate the functional states of promoters, we leveraged epigenetic data from this HAEC panel^16^ including chromatin accessibility from ATAC-seq (assay for transposase-accessible chromatin followed by sequencing)^21^ and ChIP-seq (chromatin immunoprecipitation followed by sequencing) data for the histone modification H3K27ac (histone 3 lysine 27 acetylation) that marks active regulatory elements including promoters^22^. Consistent with AF-DSTs being driven by alternative promoters, chromatin accessibility and H3K27ac were enriched at transcription start sites (TSS) of alternative first exons at both promoter types (**Figure 2B**). H3K27ac peaks widen with IL1β treatment at P_inducible_, indicating a recruitment of transcriptional machinery and widening of the nucleosome free region. The pattern of chromatin accessibility at P_inducible_ and P_basal_ is different H3K27ac, with P_basal_ having more accessibility in both IL1β and control treatments than P_inducible_ (**Supp Figure 2A**).

To see if the abundance of H3K27ac or chromatin accessibility changed at the P_inducible_ and P_basal_ upon IL1β exposure, we calculated the ratio of epigenetic sequence reads in IL1β relative to control (**Figure 2A**). Overall, there is more H3K27ac with IL1β treatment for both P_inducible_ and P_basal_, yet comparing ratios of P_inducible_ to P_basal_, there is a greater induction at P_inducible_ for the majority of AF-DSGs (**Figure 2D**). Across all AF-DSGs, 56 of 106 AF promoter pairs had greater H3K27ac induction with IL1β compared to control cells (t-test p < 0.05) (**Figure 2C**). Contrarily, chromatin accessibility at AF promoters were less dynamic than H3K27ac (**Figure 2C, Supp Figure 2A**). This supports a model whereby distinct promoters select first exon usage of the same genes based on chromatin dynamics in response to IL1β-induced signaling.

### P_inducible_ and P_basal_ sequences are enriched for distinct TF motifs

To gain insight into the DNA sequences and corresponding TFs that coordinate alternative promoter selection, we performed motif enrichment analyses using the respective DNA sequences with promoters defined as 1kb upstream to 0.5 kb downstream of each TSS. We found that ETS, KLF, NFY, and SP1 motifs were enriched in both P_inducible_ and P_basal_ relative to the genome at large (**Supp Figure 2B**). This is consistent with our work and others demonstrating that the ETS motif is a major determinant of regulatory function in ECs^23^. The ETS motif is bound by ETS family proteins (e.g., ERG, FLI1, ETS1/2, ETV2/6, ELK3) that regulate EC development and homeostasis.

Next, to gain insight into the transcription factors regulating use of the alternative promoter sets, we searched for enriched DNA motifs in the P_inducible_ and P_basal_ sequences. Several significant enrichments were observed (**Figure 2E, Supp. Table 3**). Most notably, the Jun-AP1 and E2F were enriched in P_basal_ sequences relative to P_inducible_ sequences (**Figure 2E**). In contrast, a homeobox motif, Glucocorticoid Response Element (GRE), and another E2F motif were enriched in P_inducible_ sequences relative to the P_basal_ set. Interestingly, we find different variants of E2F motifs enriched in P_basal_ and P_inducible_. The P_basal_ E2F motif is described to bind E2F1-3 proteins, which typically activate transcription during G1/S transition^24^. In contrast, the P_inducible_ enriched E2F motif is typically bound by E2F7 that is primarily classified as a repressor^25^ (**Figure 2E**). Also, enrichment of the GRE at P_inducible_ sites is consistent with the known roles of nuclear receptors regulating transcription in response to inflammation^26^. Together, these results support that distinct sets of transcription factors orchestrate alternat promoter usage during the HAEC inflammatory response.

### Transcription factors ERG and RELA tune alternative promoters

Next, to evaluate alternative promoters based on their epigenetic signatures across the HAEC lines, we extended the analysis from H3K27ac and chromatin accessibility to include chromatin binding data of two important TFs: RELA (the 65 kDa subunit of the NFkB complex) and the EC lineage-determining TF ERG (Ets-related gene). From ChIP-seq data for RELA and ERG that were collected from the same EC cohort^16^, we confirmed that both TFs bind at P_basal_ and P_inducible_ (**Supp Figure 3A-B**). Both the RELA and ERG binding peaks are wider at P_inducible_ than P_basal_ with IL1β treatment compared to control (**Supp Figure 3A-B**). This is consistent with the widening of the H3K27ac peak (**Figure 2D**) at P_inducible_ with IL1β treatment, and the model whereby chromatin remodeling primes P_inducible_ for increased transcriptional machinery upon IL1β treatment.

**Figure 3.**
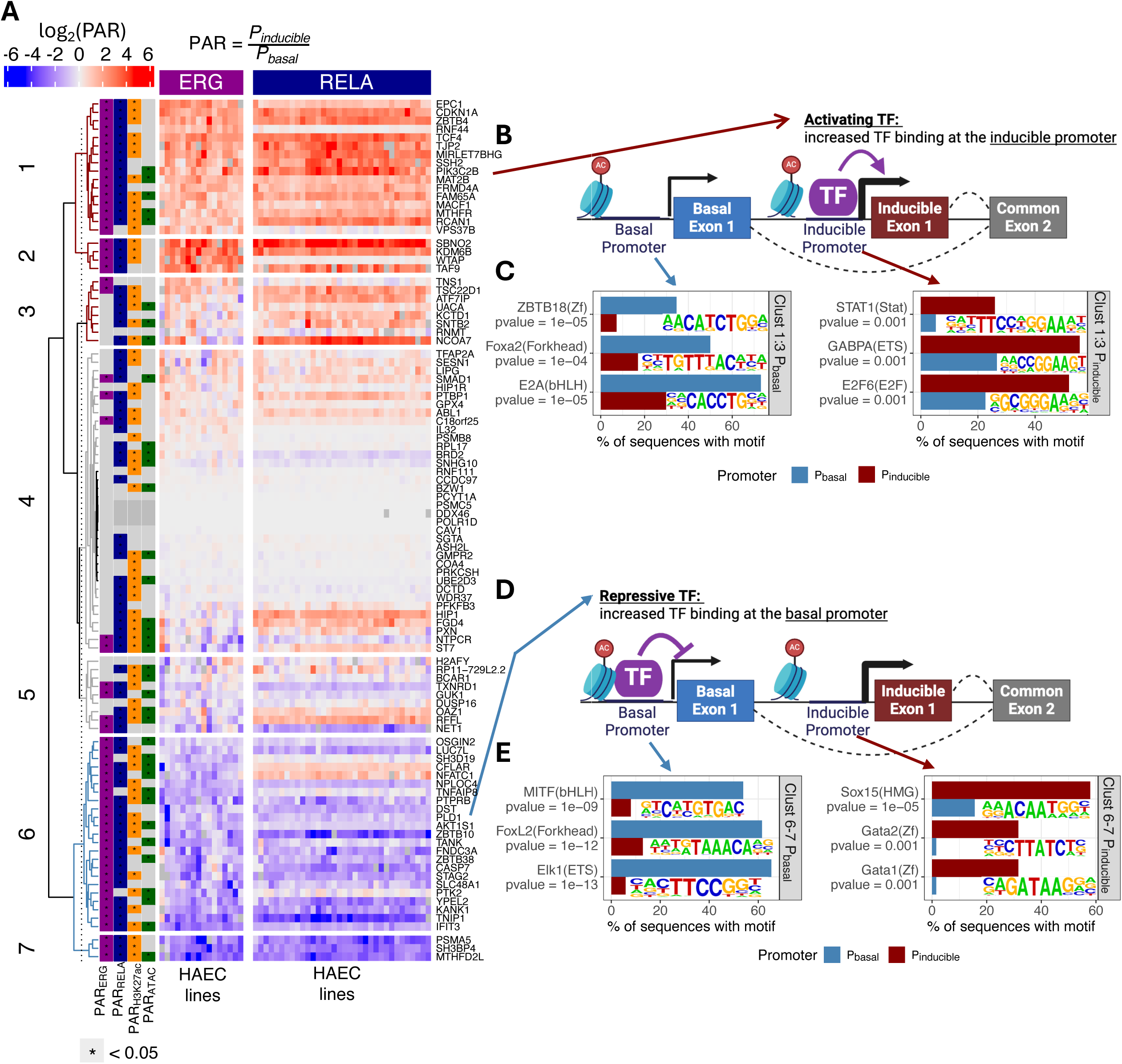
Transcription factors ERG and RELA mediate alternative promoter usage. A. PAR_ERG_ (n = 15) and PAR_RELA_ (n = 34) for 121 AF-DSGs. The log2(PAR_TF_) is plotted for visualization, log2(PAR_TF_) > 0 is activating, and <0 is repressive. The dendrogram was cut to reflect three main branches: ERG and RELA activating clusters 1-3 (red), ERG and RELA repressing clusters 6-7 (blue) and ERG and RELA ambiguous clusters 4-5 (gray). B. Diagram of the definition of an activating TF where more TF binding at P_inducible_ is correlated with more mRNA_inducible_. C. Motif enrichment in ERG and RELA activated clusters 1-3 for P_basal_ (left) P_inducible_ (right). D. Diagram of the definition of an activating TF where more TF binding at P_inducible_ is correlated with more mRNA_inducible_. E. Motif enrichment in ERG and RELA repressed clusters 6-7 for P_basal_ (left) P_inducible_ (right).

To quantitatively compare epigenetic signatures between promoters, we defined a **Promoter Activity Ratio (PAR)** metric. The PAR is the ratio of normalized sequenced reads at the P_inducible_ relative to the P_basal_ for each AF-DSG. **Figure 3A** shows PARs for ERG and RELA for each AF-DSG (rows) and HAEC line (columns). Genes with statistically significant PARs are indicated by the leftmost columns (paired t-test p < 0.05) for ERG, RELA, H3K27ac, and ATAC-seq (**Figure 3A**). Consistent with previous reports, these data indicate that NFkB and *ERG* co-localize at promoters^23^(**Supp Figure 3B**). Unsurprisingly, 50 promoter sets have a significant PAR for both ERG and RELA (**Figure 3A, Supp Figure 3D**).

The heatmap of PARs for ERG and RELA in **Figure 3A** showed a distinctive trend: promoters for genes in clusters 1-3 have TF binding trends that *<u>positively correlate</u>* with splicing patterns, whereas gene promoters in clusters 6-7 exhibit TF binding patterns that *<u>negatively correlate</u>* with splicing patterns. Log_2_(PARs) greater than 0 in clusters 1-3 mean that more ERG and RELA binding were observed at P_inducible_ than P_basal_. In simplistic terms, these data are consistent with ERG and RELA functioning as activating TFs for these genes (**Figure 3B**). Conversely, Log_2_(PARs) less than 0, as in clusters 6-7, indicate that more ERG and RELA binding were observed at P_basal_ relative to P_inducible._ In simple terms, this indicates that ERG and RELA function as repressors at P_basal_ (**Figure 3D**) because there is more promoter activity yielding AF splicing from P_inducible_. Importantly, the labels of activating and repressive are relative insofar as the same regulatory element could be bound by a repressive complex under one treatment condition and switch to become activating in the other condition. These terms are used to define groups and are not definitive. The other clusters, clusters 4 and 5, are mostly made up of promoter sets with contradictory PAR_TF_ directions for ERG and RELA, or lack of regulation by one or both TFs (**Figure 3A**).

### Motif enrichment of ERG and RELA regulated alternative promoters

To further characterize mechanisms controlling AF promoters, we performed motif enrichment for promoter sets in the gene clusters from **Figure 3A**. For clusters 1-3 (ERG and RELA ‘activated’, **Figure 3B**), the P_inducible_ sequences were enriched for STAT1, ETS, and E2F6 motifs relative to P_basal_ sequences (**Figure 3C**). Interestingly, E2F6 is often characterized as a repressor and may be binding at P_inducible_ sequences to repress activity under basal conditions^27^. Conversely, STAT1 is known to be activated by inflammatory signaling cascades and to be a binding partner of RELA^28^. In clusters 1-3 P_basal_ sequences we found the ZBTB18 motif to be enriched (**Figure 3C**). ZBTB18 is a transcriptional repressor and is known to reduce chromatin accessibility^29^ and is likely to be active under inflammatory conditions to repress transcription from P_basal_. We also identified FOXA2 and E2A motifs to be enriched in clusters 1-3 P_basal_ (**Figure 3C**). FOXA2 is a known lineage-determining factor in EC development^30^, while the role of E2A is lesser known in ECs.

Next, we identified enriched motifs in ERG and RELA repressed promoter sets (clusters 6-7) (**Figure 3D-E**). First, in P_basal_ sequences we found the MITF and FOXL2 motifs to be enriched relative to P_inducible_ (**Figure 3E**). There is evidence MITF regulates vascular endothelial growth factor^31^, but the role of FOXL2 in endothelial cells is lesser known. Conversely, in the P_inducible_ sequences (where there is less ERG and RELA binding) had GATA and SOX enriched motifs (**Figure 3E**). We previously found that SOX and GATA motifs are enriched in HAEC enhancers and were less correlated with increased chromatin accessibility than the ETS motif^23^. This is consistent with SOX and GATA acting as repressors, and these new findings suggest a novel role for SOX and GATA as repressors of inducible sites under basal conditions.

Perhaps most interesting, we identified that the ETS motif is enriched in P_inducible_ for clusters 1-3, and P_basal_ for clusters 6 and 7 (**Figure 3C, E**). Surprisingly, we did not identify any enrichment of the NFkB motif in alternative promoter sets. Therefore, we conclude that RELA does not bind chromatin through direct DNA motifs at these sites, but rather that RELA binds to complexes that are tethered to DNA by other TFs such as ERG^27^.

### sQTL mapping in HAECs

Given the extent of splicing differences we observe in ECs, we hypothesized that genetic polymorphisms in humans tune each individual’s splicing profiles with cell-specificity. We performed sQTL mapping in the control and IL1β -treated HAEC datasets (**Figure 4A**). Focusing on cis-sQTLs (within 1 Mb of the splice junctions) and using a 5% locus-wide false discovery rate, we identified 3,016 and 2,858 sQTLs in control and IL1β treatments, respectively. Of these sQTLs, 619 (12%) were significant in both cell conditions (**Figure 4B**). The 5,734 total sQTLs correspond to AS of 5,255 introns in 2,947 genes (sGenes) and 4,854 SNPs (sSNPs). Only 13% of HAEC sQTLs were already present in GTEx, the largest collection of sQTLs available with at most 6.8% sharing with any single tissue^32^ (**Figure 4B**). This again confirms that AS is cell-type specific and underscores the value in our novel HAEC dataset.

**Figure 4.**
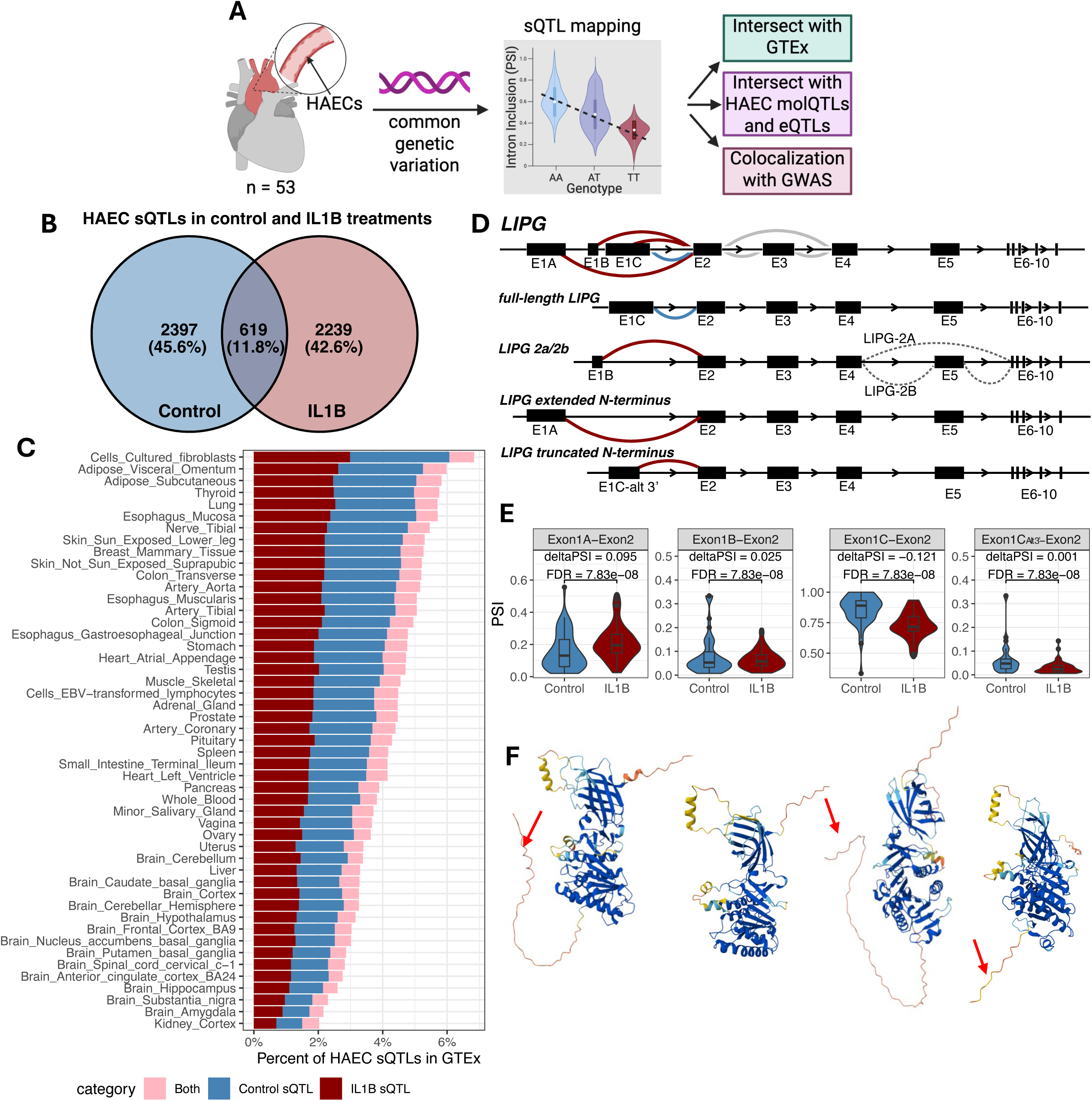
Splicing quantitative trait loci in HAECs. A. Overview of the sQTL mapping methods. B. Venn diagram of sQTLs in Control and IL1B treatments C. HAEC sQTL overlap with GTEx tissues. Bars are colored by whether the HAEC sQTL was significant in Control (blue), IL1B (red), or both treatments (pink). D. LIPG differential splicing, red splices indicate an increase with IL1B, blue splices a decrease with IL1B, and gray splices are regulated by an sQTL. LIPG differential splicing with IL1B can produce full-length LIPG, LIPG 2a/2b, LIPG with extended N-terminus, or LIPG with truncated N-terminus, the last two of which include previously undescribed first exons. LIPG 2a is made from the E1B first exon and includes exon 5, while LIPG 2b starts with E1B and excludes exon 5. E. LIPG AF differential splicing between IL1B and Control treatment in HAECs (n = 53), FDR indicates benjamini-hochberg corrected FDR for differential splicing of the cluster by leafcutter. F. LIPG predicted protein structures (AlphaFold ^71^)with the canonical amino acid sequence from full-length LIPG containing the signal peptide domain indicated by the red arrow (left), LIPG-2A (middle left) which contains no signal peptide domain, LIPG with extended N-terminus with the extra 34 amino acids at the N-terminus marked by the red arrow (middle right), and LIPG with truncated N-terminus with red arrow pointing to the truncated signal peptide domain (right).

We were curious if the sQTLs were enriched in other molecular QTLs (molQTLs) or eQTLs that were previously reported using this HAEC cohort^16^. The available molQTLs included QTLs for ERG binding, RELA binding, chromatin accessibility, and histone QTLs for H3K27ac. Of these traits, sSNPs were most likely to also have significant eQTLs, followed by histone H3K27ac QTLs (**Supp Figure 4A**).

### LIPG: an example of an EC -specific sQTL

Lipase G, Endothelial-type (LIPG) is an example of a gene that is regulated by both IL1β and genetic variation (**Figure 4D**). *LIPG* is differentially spliced with IL1β treatment (FDR = 7.38 × 10^−8^) at its 5’ end and can produce four possible first exons (**Figure 4D-E**). To our knowledge, two alternative first exons have been described for *LIPG*^33^: exon 1A that produces full-length protein (LIPG-1a), and exon 1b, which is upstream and utilizes an internal start codon thereby producing a truncated first exon (LIPG-2a/b) (**Figure 4D**). We identify this same alternative first exon 1b and confirm that it is upregulated with IL1β treatment ^33^ (**Figure 4E**). We identified two other novel first exons for *LIPG*: exon 1c, which is further upstream than exon 1b and encodes an additional 34 amino acids, and exon 1d that uses the same TSS as the canonical exon 1a but has an alternative 3’ end that skips the protein-coding nucleotides in exon 1c, removing 32 amino acids (**Figure 4E**). The predicted protein structures for these *LIPG* transcripts differ in their amino terminal signal peptide domains: LIPG-2a does not have the signal peptide domain, LIPG-truncated contains a truncated signal peptide, and LIPG-extended has an extra 34 amino acids in that domain (**Figure 4F**).

*LIPG* is also regulated by genetics as rs9944692 is an sQTL for two intron inclusions in LIPG downstream from the alternative start sites (**Supp Figure 4B-C**). This sQTL is not published in GTEx and given that *LIPG* is an endothelial-specific gene it is unsurprising that its discovery required a single cell type culture. Exon 3 encodes amino acids 93-153 which are proximal to the lipid binding domain of LIPG, and its omission may have structural effects for the protein or affect its catalytic activity ^33^ (**Supp Figure 4E**). Interestingly, the sQTL identified does not solely regulate the skipping of exon 3, but also a potential truncation event. The rs59944692-AA genotype produces more junctions between exons 2 and 3 and fewer between exons 3 and 4, suggesting that the rs59944692-AA genotype is likely to produce transcripts that end with exon 3 (**Supp Figure 4B-C**). If the transcript were to be truncated after exon 3, the protein would be missing more than half of its amino acids on the C-terminal end (**Supp Figure 4E**). The sSNP rs59944692 is also an ERG binding QTL^16^, and rs59944692-AA produces more ERG binding than rs59944692-AG (**Supp Figure 4F**). This ERG binding site also has characteristic enhancer marks and may be a previously undescribed enhancer for *LIPG* splicing (**Supp Figure 4G**). This *LIPG* splicing enhancer is also nearby an eQTL sSNP for *LIPG* (**Supp Figure 4G**). This enhancer site is downstream of the *LIPG* locus and contains 5 potential enhancer-like structures containing sQTL and eQTL SNPs for LIPG (**Supp Figure 4G**). Since these signals do not colocalize at the same genetic variant, this enhancer region is likely to have two functions, to regulate splicing and expression of the *LIPG* locus.

### sQTLs colocalize with GWAS signals

Lastly, we aimed to identify how common genetic variants may affect complex disease through AS. To do so, we used genetic colocalization analysis to statistically identify sQTLs that share genetic association signals with disease-modifying SNPs from GWAS. Several vascular biology associated traits were considered including total cholesterol levels, low-density lipoprotein levels (LDLC), high-density lipoprotein levels (HDLC), diastolic and systolic blood pressure (BP), pulmonary embolism (PE), deep vein thrombosis (DVT), and coronary artery disease (CAD)^34,35^. We found that LDLC and Diastolic BP had the greatest number of colocalizing signals with HAEC sQTLs (13 out of 53, corresponding to 9 and 6 sGenes, respectively) (**Figure 5A**, **Table 1**). Splices in the four genes *M6PR, PSORS1C1, PROCR,* and *RBM23* colocalize with the most GWAS traits (**Figure 5A**).

**Figure 5.**
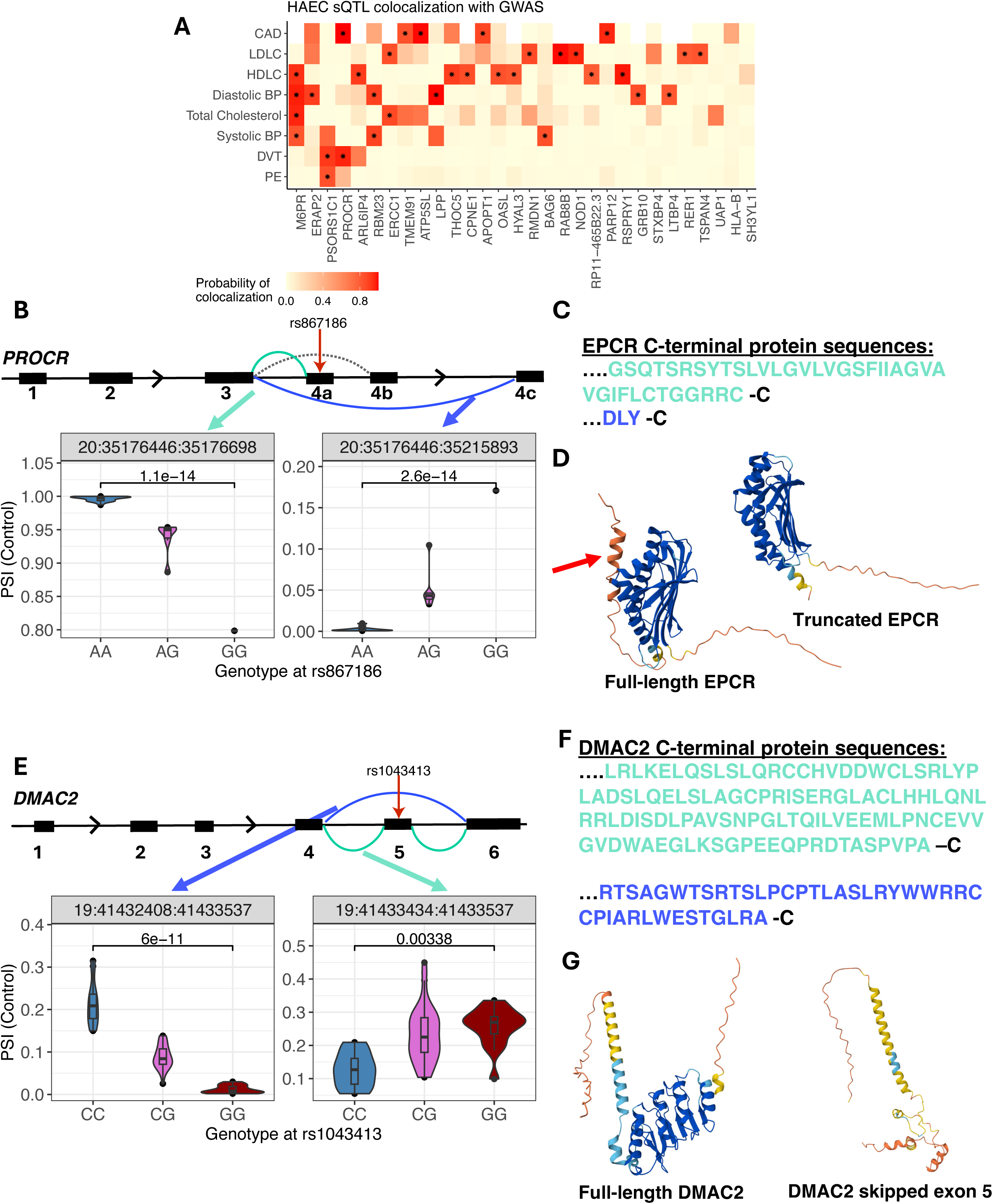
HAEC sQTLs colocalize with GWAS signals for cardiovascular traits and disease. A. Heatmap of posterior probability of colocalization of sQTLs with GWAS traits by gene symbol and trait. Significant colocalizations are indicated by asterisk (PP > 0.8). B. PROCR locus with sQTL splices highlighted from exon 3-4a (canonical) and exon3-4c (novel). PROCR AS by genotype at rs867186 in Control treated HAECs, FDR represents locus level FDR for the sQTL. C. EPCR protein sequences for *PROCR* exon 4a (full-length, green) and exon 4c (truncated, blue). D. EPCR predicted protein structures (AlphaFold^71^) with *PROCR* exon 4a (full-length, left) and exon 4c (truncated, right) as the last exon. The red arrow indicates the transmembrane domain that is missing from truncated EPCR. E. DMAC2 locus with sQTL splices highlighted from exons 4-5, 5-6, and 4-6. DMAC2 AS by genotype at rs1043413 in Control treated HAECs, FDR represents locus level FDR for the sQTL. F. DMAC2 protein sequences for *DMAC2* containing exon 5 (full-length, green) and skipping exon 5 (blue). G. DMAC2 predicted protein structures (AlphaFold ^71^) with containing exon 5 (full-length, left) and skipping exon 5 (right).

### AS of PROCR is associated with CAD and DVT

The *PROCR* gene encodes the Endothelial cell protein C receptor (EPCR), a glycoprotein that exists on the luminal surface of large vessels and in plasma in its soluble form (sEPCR)^36^. EPCR is solubilized and released into circulation upon proteolytic cleavage in a domain encoded in exon 4a^37^ or by AS in exon 4b that results in deletion of the transmembrane domain ^36^. The rs867186 A/G polymorphism causes a Ser219Gly non-synonymous amino acid substitution. The A allele is associated with CAD^35^, while the G allele is associated with venous thromboembolism (VTE) ^38^, and increased sEPCR^36^. rs867186 is an sSNP for *PROCR* for splices between exons 3-4a and 3-4c (**Figure 5B**). The A allele associates almost exclusively with transcripts that utilize the canonical last exon, 4a, while the G allele significantly associates with increased of exon 3-4c splices (**Figure 5B, Supp Figure 5A**). We confirm that the A allele results in more splicing from exon3-4a but did not identify an association between genotype at rs867186 and exon-4b. Interestingly, rs867186 is also an eQTL for *PROCR,* with the A allele producing more expression (**Supp Figure 5B**). The novel *PROCR* exon 4c contains only 6 protein-coding amino acids, a stop codon, and 3’ UTR sequence (**Figure 5C**). Although the *PROCR* sQTL is in GTEx (**Supp Figure 5C**) to our knowledge the function of the alternative 4c last exon is not known. Thus, we validated the existence of the 4c exon using PCR from poly-A selected cDNA in an EC line heterozygous for rs867186 and confirmed that there is *PROCR* mRNA made containing this novel 4c exon (**Supp Figure 5D**).

The predicted protein structure of exon-4c containing EPCR is truncated and missing the transmembrane domain (**Figure 5D**, indicated by the red arrow), similar to sEPCR (**Supp Figure 5H**). Based on sequence and predicted structure, we can hypothesize that the inclusion of exon 4c results in a decrease in full-length, membrane bound EPCR and more sEPCR, but validation at the protein level will be required for definitive proof of this hypothesis. Using colocalization analysis we found that the *PROCR* sQTL shares a genetic association with CAD ^35^ (PP = 1, **Table 1, Supp Figure 5C-D**) and deep vein thrombosis ^32^ (DVT) (PP = 0.96, **Table 1, Supp Figure 5 E**). To our knowledge this is the first time that *PROCR* AS and CAD or DVT disease-status have been evaluated together. This analysis presents a unique opportunity to explore the function of the novel, truncated exon-4c containing *PROCR* transcript in cardiovascular disease.

### AS of *ATP5SL/DMAC2* is associated with CAD

Another sQTL that colocalizes with CAD is for the sGene Distal Membrane Arm Assembly Component 2 (*DMAC2,* sometimes referred to as *ATP5SL*) (PP = 0.99). DMAC2 is required for assembly of complex I in the mitochondria^39^. The *DMAC2* sQTL identified regulates skipping of exon 5 (**Figure 5E, Supp Figure 6A**). The sSNP, rs1403413, is in exon 5 and disrupts a splicing enhancer sequence^40^. We verified that this splicing event occurs with PCR and confirm that two transcripts are made, one with and one without exon 5 (**Supp Figure 6D**). This sQTL is present in GTEx in several tissues, including the aorta (**Supp Figure 6E**). However, to our knowledge the functional consequence of the loss of exon 5 on protein function has not been described. When exon 5 is omitted from the transcript, the protein is 99 amino acids shorter (**Figure 5F**). The predicted protein structure of the canonical protein sequence is markedly different—an indication that omission of exon 5 is likely to alter protein function, namely ATP synthesis (**Figure 5F**).

The sSNP rs1043413 is also a GWAS SNP for CAD^35^ (p = 3.66e^−8^) (**Table 1**). rs1043413 is the lead SNP for the sQTL for *DMAC2* (**Figure 5F**) but is not the lead SNP for CAD, although it is in high LD (> 0.8) with the lead SNP rs4574 (**Figure 5G**). rs4574 was tested for association with the splice but was not significant (**Figure 5F**). This suggests that there are two signals at the *DMAC2* locus and that rs4574 may regulate of another gene in the region, or that the signal from rs4574 and splicing of *DMAC2* are independently associated with CAD.

## Discussion

This study is to our knowledge the most comprehensive analysis of transcriptome-wide RNA splicing in HAECs. Using a genetically diverse panel of 53 HAEC lines, 1,224 transcripts were significantly differentially spliced upon IL1β exposure (**Figure 1**) and splicing of 2,947 sGenes varied significantly as a function of genetic variation (**Figure 4**). Among DSGs, we identified enrichment in genes with known function in metabolic reprogramming and stress response pathways (**Figure 1D**). We identified an abundance of AF exon usage with IL1β exposure, which in conjunction with epigenetics data were confirmed to be driven by alternative gene promoter activities (**Figure 2)**. This led to molecular characterization of a putative novel isoform of PFKFB3 (**Figure 1E**), whose protein function warrants further investigation. Next, we leveraged genomic sequences at AF promoters with distinct epigenetic profiles (**Figures 2-3**) to identify enriched motifs for TF families that likely direct alternative first exon selection in the HAEC transcriptional response to IL1β. Lastly, we linked quantitative splicing rates across the 53 HAEC lines to genetic variation using sQTL mapping and identified tens of human loci that colocalized with disease risk signals in GWAS. Paramount among these was a novel splice and 3’ sequence for the CAD-associated gene *PROCR*, which encodes EPCR. Taken together, this study provides the vascular biology and human genetics communities with an extensive resource to further insight into endothelial dysfunction and protein diversity. The major findings are discussed in turn below.

DSG by IL1β were enriched in metabolic reprograming and stress response pathways (**Figure 1**). These pathways are known to contribute to endothelial dysfunction^2,41^ and are demonstrated to initiate CAD, hypertension and diabetes^2^. IL1β activates the NLRP3 inflammasome, which in turn produces more reactive oxygen species (ROS) and exacerbates oxidative stress^42^. Of the DSGs in the oxidative stress pathway (**Supp. Table 2**), only RCAN1 and NCOA7 were characterized previously as DSGs under inflammatory conditions^43,44^. RCAN1 is differentially expressed under hypoxic conditions in a HIF1a dependent manner, and the RCAN1 AS isoform regulates VEGFR2 signaling in ECs^45,46^. It was shown that RCAN1 splicing is mediated by an alternative promoter that is activated by the NFkB complex^44^, which is consistent with our observations in this study (**Figure 3C**). While the known involvement of oxidative stress and metabolic reprogramming are not new, the specific splices identified in this study are >95% specific when comparing ECs to macrophages (**Supp.** Fig 1E), indicating that the majority of events detected are cell type specific.

We identified a novel IL1β -regulated AF for the glycolytic enzyme PFKFB3 (**Figure 1E**). The canonical PFKFB3 has demonstrated roles in angiogenesis, sprouting, and vascular development^47–50^. It also promotes tumorigenesis through the Warburg effect whereby cancer cells increase survival in hypoxic environments. Small molecule inhibitors of PFKFB3 have been explored as anti-cancer therapies with some promising results in pre-clinical animal models^51,52^. Our data identify for the first time that an alternatively spliced exon exists upstream of the PFKFB3 substrate binding domain that catalyzes synthesis of fructose-2,6-bisphosphate^53^. Although some AS of PFKFB3 has been reported near the 3’ end of the gene^54^, our novel ‘isoform 2’ is not described in the literature. Studies of PFKFB3 generally refer to isoform 1, albeit most reagents (e.g., antibodies, interfering RNAs etc) would target both isoforms given the sequence identity after exon 1. We suggest that the alternative first exons of PFKFB3 may play a role in regulation of glycolysis and that in response to IL1β the dominant form of PFKFB3, isoform 1, is favored.

We also report that AS of *DMAC2*, another metabolic gene, is regulated by AS (**Figure 6E-G**). DMAC2 is an essential component of complex I formation in the mitochondria ^39^, which is conserved across all three domains of life. We found that AS of *DMAC2* by genotype at rs1043413 is significantly colocalized with CAD (**Table 1**). Complex I deficiencies have been linked to diverse clinical phenotypes, including cardiomyopathies, neurodegenerative disorders, liver disease, CAD, and stroke^55^.

During ischemia, complex I undergoes a switch to a dormant state in the heart and brain that involves a conformational change in the complex^56^. Skipping of *DMAC2* exon 5 produces a frameshift in the coding sequence, and a drastic change to the predicted protein structure (**Figure 5G**). Thus, we suggest that AS of *DMAC2* may be a potential new target for this area of research.

We identified 340 AF-DSTs and that AF splices are significantly enriched in this analysis (**Figure 1B**). Furthermore, AF-DSGs represent a significant proportion of DSGs in enriched biological pathways and are likely to result from alternative promoter usage based on in H3K27ac (**Figure 2B**), and binding of ERG and RELA (**Figure 3A, CA).** Both ERG and RELA are known regulators of gene expression in response to inflammation and it is unsurprising that they modulate gene expression in response specifically to IL1β treatment^16,23,57,58^. Here we demonstrate for the first time that ERG and RELA are master regulators of AS via regulation of alternative promoters (**Figure 3**). In endothelial cells, ERG is lineage-determining^59^ and to be repressive of endothelial dysfunction, including EndMT^60^ which is a process correlated with the progression of atherosclerosis^61^. The role of ERG in AS has been previously unappreciated. ERG knockdown induces pro-inflammatory gene expression and sensitizes cells to IL1β treatment and to EndMT-signaling^23,62^. We can conclude from this that ERG plays a role in the splicing-level response to IL1β stimulus and points to another potential mechanism by which ERG maintains homeostasis in ECs under inflammatory conditions.

In contrast to ERG, The NFkB complex is known to regulate AS in several systems. For example, the RELA subunit of NFkB binds splicing regulator DDX17 and recruits it to target exons and coordinate 3D chromatin remodeling and AS^63,64^. Given NFkB’s wide range of functions and target genes, it is unsurprising that it has a global effect on the AS landscape in response to inflammatory IL1β. The finding that the ETS motif is enriched in alternative promoters at promoters where both ERG and RELA increase binding (**Figure 3C, E**) supports a model whereby ERG directly binds DNA and tethers NFkB to chromatin. Further investigation into binding partners that regulate AS alongside NFkB in other cell types is warranted, as ERG is an EC-specific TF and NFkB has a wide breadth of influence over transcription across cell types and tissues.

Perhaps the most exciting finding in our study is the AS event we identify for the *PROCR* locus (**Figure 5B**) which significantly colocalizes with the highly reproduced genetic risk for CAD and VTE (**Figure 5C-D**, **Table 1**). The sSNP for *PROCR* rs867186-G allele is a known risk allele for venous thromboembolism^38^ and increased sEPCR^65^, while the major allele rs867186-A is associated with higher risk of CAD^35^. EPCR is an incredibly important protein in endothelial homeostasis and response to inflammatory signaling. Membrane-bound EPCR activates Protein C and is responsible for signaling cascades involved in maintaining vascular barrier integrity, local anticoagulatory function, and reducing local inflammation^66^. sEPCR can still bind protein C, but it does not elicit anti-coagulatory signaling in ECs since it is not tethered to the cell. Instead, sEPCR sequesters protein C and fails to provide local anti-inflammatory and anti-coagulatory effects^37,67^. Increased levels of sEPCR are associated with renal impairment^68^, and severe malaria^69^ and can be used as a biomarker for endothelial dysfunction^67^. The novel AL exon we identify for *PROCR* is likely to increase sECPR as it skips the exon that encodes the transmembrane domain, but further studies into the truncated EPCR protein structure will be necessary.

In conclusion, we present a comprehensive analysis of AS in human ECs both by IL1β and common genetic variation. The findings presented here support that ECs express numerous novel transcripts that are relevant to cardiovascular disease. These data will serve as a resource to the research community to accelerate the discovery of new targets for cardiovascular disease.

## Methods

RNAseq, ChIPseq, and ATACseq next gen sequencing data from Stolze et al., 2020 are publicly available at NCBI GEO database with the accession numbers GSE30169 and GSE139377. Monocyte-derived macrophage RNAseq data was retrieved from the NCBI GEO database with the accession number GSE147310.

GTEx sQTLs: The data used for the analyses described in this manuscript were obtained from the GTEx Portal on 10/29/24.

Custom scripts will be available upon publication on GitHub: https://github.com/akgolebiewski/Golebiewski-splicing-2025

### Wet lab and cell culture

#### HAEC culture

Human aortic endothelial cells were cultured in M199 with 20% FBS (Cytiva, HyClone), ECGS (ThermoFisher #354006), Heparin, Amphotercin B (Fungizone, ABM #G274), Penicillin/Streptomycin (Gibco #15070063), and NaPr. All cells were cultured on gelatinized, tissue-culture treated plates. HAECs were used at passage six or lower to maintain native profiles.

#### Library preparation and sequencing

Sequence libraries for RNAseq, ChIPseq, and ATACseq were prepared as previously described (Hogan et al., 2017) and stated in Stolze et al, 2020.

#### cDNA preparation and PCR

RNA was extracted from HAECs using the QuickRNA Microprep kit (Zymo #1051) and then mRNA was selected using polyDT beads (Invitrogen #61002). cDNA was synthesisized using oligoDT priming and random hexamers with SuperScript III (Invitrogen # 18080051). PCR primers are cycling conditions (annealing temperatures) are listed in **Supplemental Table 5**. PCR was performed using the NEB2Next PCR Master Mix (NEB # M0541L) with the following cycling conditions: 1) 98°C for 30 seconds, 2) 98°C for 1-seconds, 3) 60-63°C for 30 seconds, 4) 72°C for 30 seconds, repeat steps 2-4 24X, 5) 72°C for 2 minutes, 6) hold at 4°C.

### Bioinformatics

#### RNAseq workflow

Sequencing data were mapped to hg38 utilizing STAR with default parameters. In our previous publication, Stolze et al., 2020, the same parameters were used to align reads to hg19 but for this analysis data were re-mapped to hg38. The resulting binary alignment file (BAM) was filtered and used for junc file creation using the Leafcutter pipeline provided here: https://github.com/davidaknowles/leafcutter. Tag directories were created using HOMER makeTagDirectory.pl and count matrices produced using AnalyzeRepeats.pl.

#### Peak calling (ATAC and ChIP)

Peaks were identified using HOMER findPeaks.pl and merged across donors using annotatePeaks.pl with the “-noadj” setting for differential analysis and “fpkm” for visualization.

#### Differential splicing

Leafcutter (https://github.com/davidaknowles/leafcutter)^17^ was utilized for differential splicing analysis. Sex and ancestry as biological variables were addressed as covariates. Replicates of RNA libraries for donors were merged before junc files were created using “bed2junc.pl” and differential splicing analysis was performed.

Significance was defined as Benjamini-Hochberg FDR < 0.05 and an effect size greater than 0.05 (−0.05 < deltaPSI > 0.05). SUPPA2^70^ was used to generate reference panels for all the splice types measured (AF, AL, SE, A3, A5, MX, RI) using the gencode annotation of hg38 (v41). Introns identified with leafcutter were matched with SUPPA2 annotated introns to identify the transcript and splice type. If there was no matching transcript from SUPPA2, annotations were imputed based on other splice events from the same gene. If there were still no matching annotations, the leafcutter assigned annotation was used.

Introns were classified as affecting protein-coding or nonsense-mediated decay (NMD) exons based on predictions from Gencode V41, or the presence of a stop codon within 50 bp of the 5’ end of the exon. We used the definition of “exon” to mean both UTR and protein-coding exons for this analysis.

#### Pathway enrichment analysis

Biological pathway enrichment analysis was performed using clusterProfiler (https://www.bioconductor.org/packages/clusterProfiler) for biological pathways (BP), molecular functions (MF), and cell compartments (CC) for all DSGs. Because DSGs can have both positive and negative effect sizes, only unique DSGs upregulated with IL1β were used for pathway analysis. DSGs were ranked by deltaPSI for the enrichment test, full results are available in Supplemental Table 2.

#### Identifying Promoter Regions

The sequencing libraries used in this analysis were sequenced with 50 bp reads and not long-read sequencing. Furthermore, our differential splicing analysis relies on reads across junctions and not across entire exons. For AF-DSGs, it was necessary to identify the 5’ end of the first exon (the TSS) inferred from the 3’ end of the first exon. To do so, we matched the 5’ end of the AS junction to the 3’ end of first exons in the Gencode V41 annotation and pulled the corresponding 5’ end of the exon. Only one TSS was used for each first exon, even if there were multiple possible 5’ ends. This is a limitation of short-read sequencing, but we did verify using the UCSC genome browser that there were RNA reads at the TSS sites we identified.

#### Motif enrichment analysis

Both DNA and RNA motif analysis was performed using Homer. DNA motif analysis was performed for promoter regions, defined as 1 kb upstream of the TSS to 0.5 kb downstream of the TSS. Motif analysis was performed with the human genome as the background (default) and with the alternative promoter as the background.

#### Calculating Promoter Activity Ratios (PAR)

We defined PAR as the ratio of sequencing reads at P_inducible_ to P_basal._ This ratio was calculated for each HAEC sample individually. To compare the effect of treatment on the PAR, we performed a T-test between the PAR in IL1B to Control for each DSG for H3K27ac and ATAC with significance defined as p < 0.05.

Similarly, we calculated the ratio of mRNA_inducible_ to mRNA_basal_ in Control and IL1B treatments and compared the two. This serves as validation of our definition of the inducible and basal promoter systems based on RNA reads.

We calculated PAR for ERG and RELA binding using the same methods, for Control and IL1B treatments, respectively. PAR was only calculated for ERG in Control treatment because ERG expression decreases significantly with IL1B treatment and this confounds the amount of ERG binding. Likewise, RELA PAR was only calculated in IL1B treatment because under basal conditions RELA is sequestered in the cytoplasm and unlikely to bind DNA frequently.

#### Accessing GWAS data

GWAS summary statistics for LDLC, HDLC, VTE, PE, total cholesterol, diastolic blood pressure, and systolic blood pressure were accessed from the Pan-UKBB study (https://pan.ukbb.broadinstitute.org/)^34^. GWAS summary statistics for CAD were accessed the GWAS catalog from van der Harst and Verweji et al. under accession number GCST005194.^35^

Summary statistics from the PanUKBB study were provided with SNP IDs in hg19. The R package “rtracklayer” was used to lift positions from the hg19 genome annotation to the hg38 genome annotation with the command “liftOver()”. Then, SNPs were matched between GWAS and sQTL data by chromosome and position in hg38.

#### GWAS Comparison

The R package “coloc” to assess colocalization of GWAS SNPs and sQTLs found in this study. This was run using crude p-values, minor allele frequencies, standard error, and sample numbers with the command “coloc.abf()”. Colocalization tests were run for each intron, restricted to SNPs that were tested in for sQTLs and GWAS. Significance was defined as posterior probability of colocalization (PP.H4) > 0.8. Tests were run separately for Control and IL1B sQTLs and significant results are summarized in Table 1.

#### Data visualization

Leafviz, from Leafcutter, was used to visualize splicing. All other data was visualized using ggplot2. Diagrams were created using BioRender.

## Additional Information

