## Supplemental figures for "Inflammation-Induced Alternative Splicing in Human Endothelial Cells Reveals Genetic Mechanisms of Cardiovascular Disease Risk"

**SUPPLEMENTAL DATA**

6 supplemental figures and figure legends, and 6 supplemental tables provided in the excel sheet.

**Supplemental Figure Legends**

**Supplemental Figure 1**

A. Volcano plot showing deltaPSI on the x-axis, and –log10(p.adjust) for DSTs between control and IL1B.
B. Flowchart of the splice type determination.
C. Histogram of DST effect sizes. Colored lines represent different splice types. Notably, AF DSTs have the largest effect sizes in both positive and negative directions.
D. Volcano plots showing differential expression between control and IL1B treatment with genes that have AF DSTs in purple (left) and SE DSTs in blue (right). Of DEGs that have DSTs, the effect size for differential expression is modest.
E. Comparison of DSTs in HAECs with IL1B treatment and monocyte-derived macrophages with LPS treatment [16].
F. Histogram of the proportion of AF-DSGs in each enriched GO term. Red dotted line indicates the overall proportion of AF-DSGs in the dataset, demonstrating an enrichment of AF-DSGs in GO terms relative to the expected proportion.
G. PFKFB3 isoform expression with 4, 8, and 24 hours of IL1B or control treatment measured with qPCR primers specific to the alternative sequences in the first exons of isoform 1 and 2 of PFKFB3. P-values reflect a paired T-test.

**Supplemental Figure 2**

A. Heatmap of PSI (RNA), and tags in promoters (H3K27ac-ChIP-seq and ATAC-seq) by HAEC donor in IL1B and control treatment at inducible and basal promoters
B. Motif enrichment in Pbasal and Pinducible sequences relative to the genome, the top 10 results are plotted. ETS (ERG) motifs are labeled with a red box.

**Supplemental Figure 3**

A. RELA binding at alternative promoters in HAECs. Histograms representing the fragment depth per bp per peak centered at the TSS.
B. ERG binding at alternative promoters in HAECs. Histograms representing the fragment depth per bp per peak centered at the TSS.
C. Upset plot showing significant PAR values for ERG, RELA, H3K27ac, and ATAC. "+" direction indicates more signal at Pinducible vs Pbasal, "-" direction indicates more signal at Pbasal vs Pinducible.

**Supplemental Figure 4**

A. sQTL overlap with eQTLs and molQTls

B. LIPG locus with AS events indicated. Differential splicing of LIPG is indicated by dark gray splices from exons 1a, 1b, and 1c to exon 2. sQTL splices are between exons 2,3 and 4.
C. LIPG splicing by genotype at rs59944692 in Control treated HAECs. FDR represents the locus level FDR for the sQTL.
D. ERG binding (ERG molecular QTL) at a distal LIPG enhancer by genotype at rs69944692. FDR = 0.44.
E. LIPG predicted protein structure from Alphafold. Red arrow indicates the site that will be modified if exon 3 is omitted, or where truncation would occur if the transcript does not include exons 4-10.
F. PCR products validating the LIPG sQTL splices. PCR primers from LIPG exon 1a to LIPG exon 6 (sequences in supplemental table 5) produce a 1026 bp product if exon 3 is included and 847 bp product if exon 3 is excluded.
G. LIPG enhancer region in HAECs with IL1B or Control treatment

**Supplemental Figure 5**

A. PROCR eQTL at rs867186 in HAECs
B. PROCR sQTL in IL1B treatment at rs867186 in HAECs
C. PROCR sQTL for GTEx Artery-Tibial tissue for rs867186
D. PROCR validation PCRs. E1-E4: PROCR exon 1-4 with expected product 1026 bp, E1-5: PROCR exon 1-5 (novel) with expected product 841 bp, NTC: no template control.
E. PROCR sQTL locus zoom
F. PROCR CAD locus zoom
G. PROCR VTE locus zoom
H. sEPCR predicted protein structure (Alphafold)

**Supplemental Figure 6**

A. ATP5SL (DMAC2) sQTL in IL1B treatment at rs1043413 in HAECs
B. ATP5SL sQTL locus zoom
C. ATP5SL CAD locus zoom
D. ATP5SL (DMAC2) GTEx Artery-Aorta sQTL for rs1043413

**SUPPLEMENTAL TABLES**

| TABLE # | SHORT DESCRIPTION |
|---|---|
| 1 | DSTs IL1$\beta$ vs control |
| 2 | Pathway enrichment (overall, IL1$\beta$ vs control) |
| 3 | Motif enrichment Pbasal vs Pinducible |
| 4 | Motif enrichment in ERG and RELA promoter sets |
| 5 | Primers used for AS validation |
| 6 | Protein sequences used for alphafold predictions |

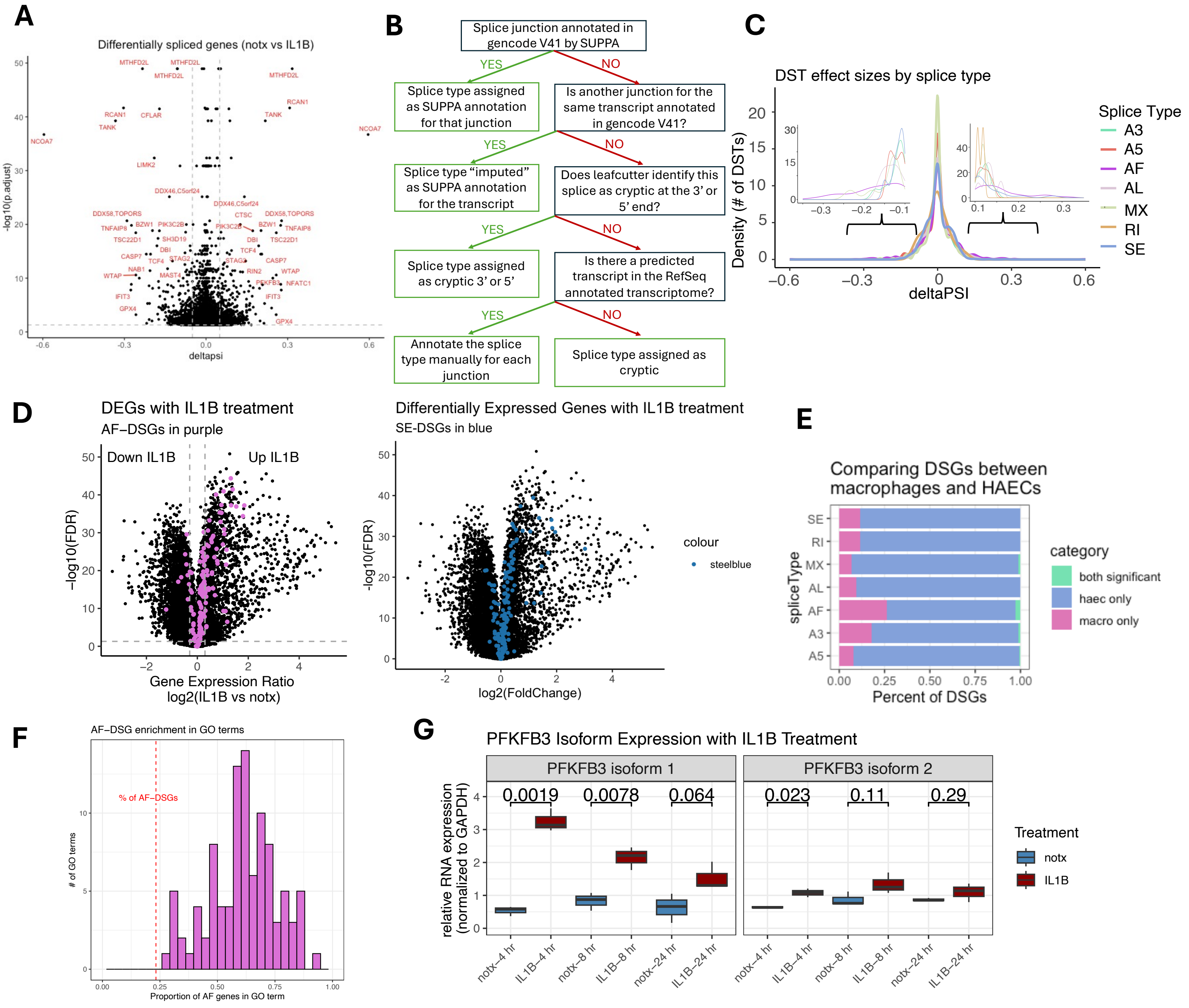


### Supplementary Figure 1

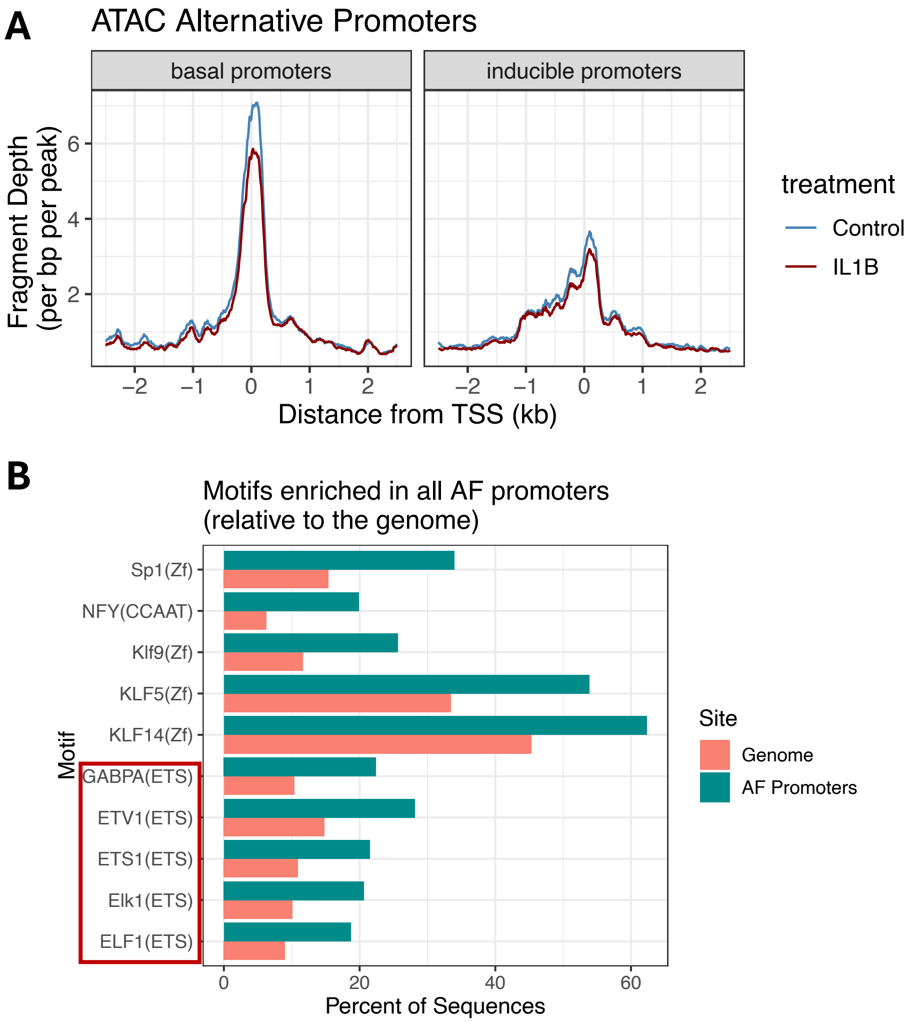


### Supplementary Figure 2

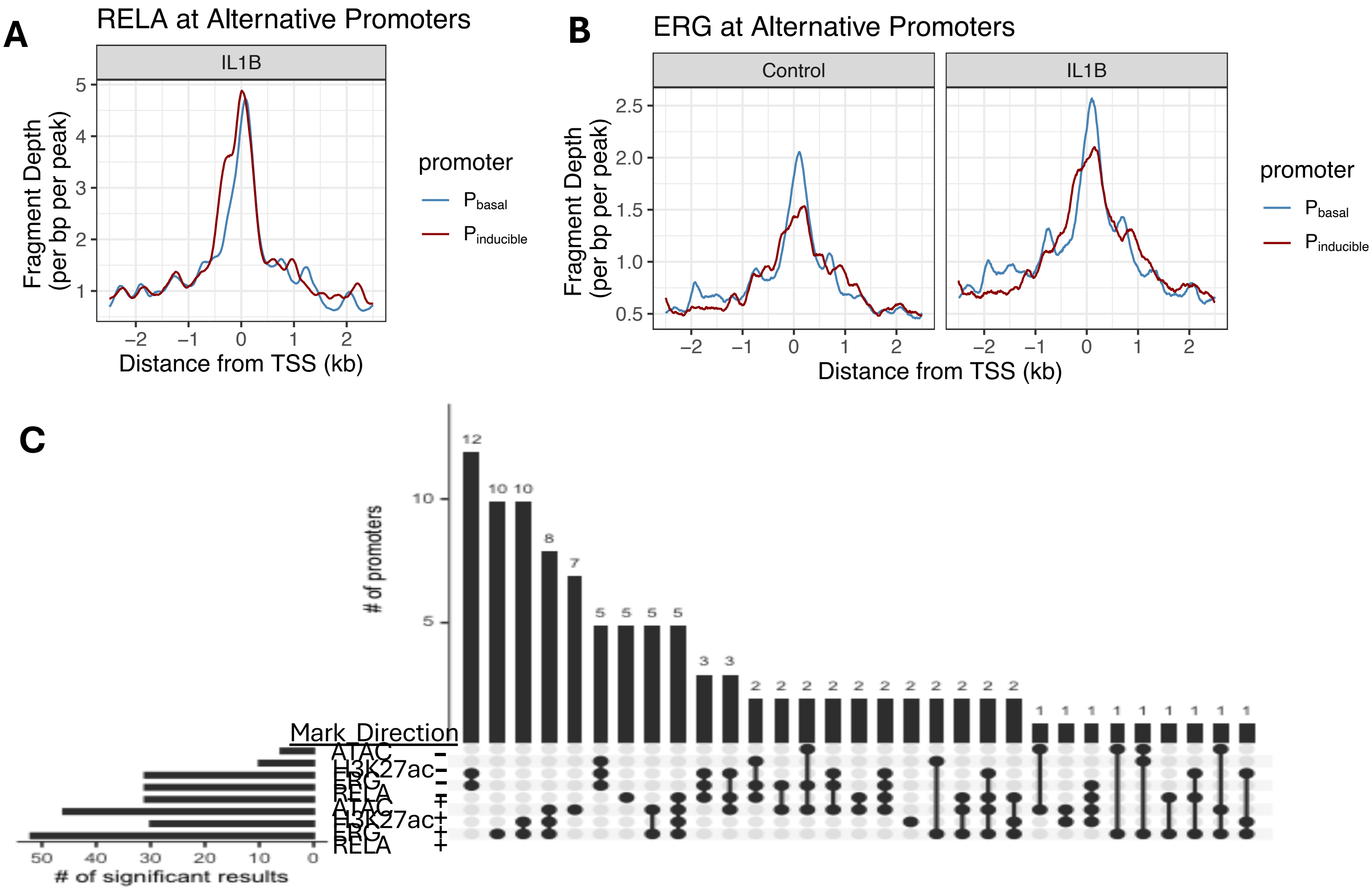


**Supplementary Figure 3**

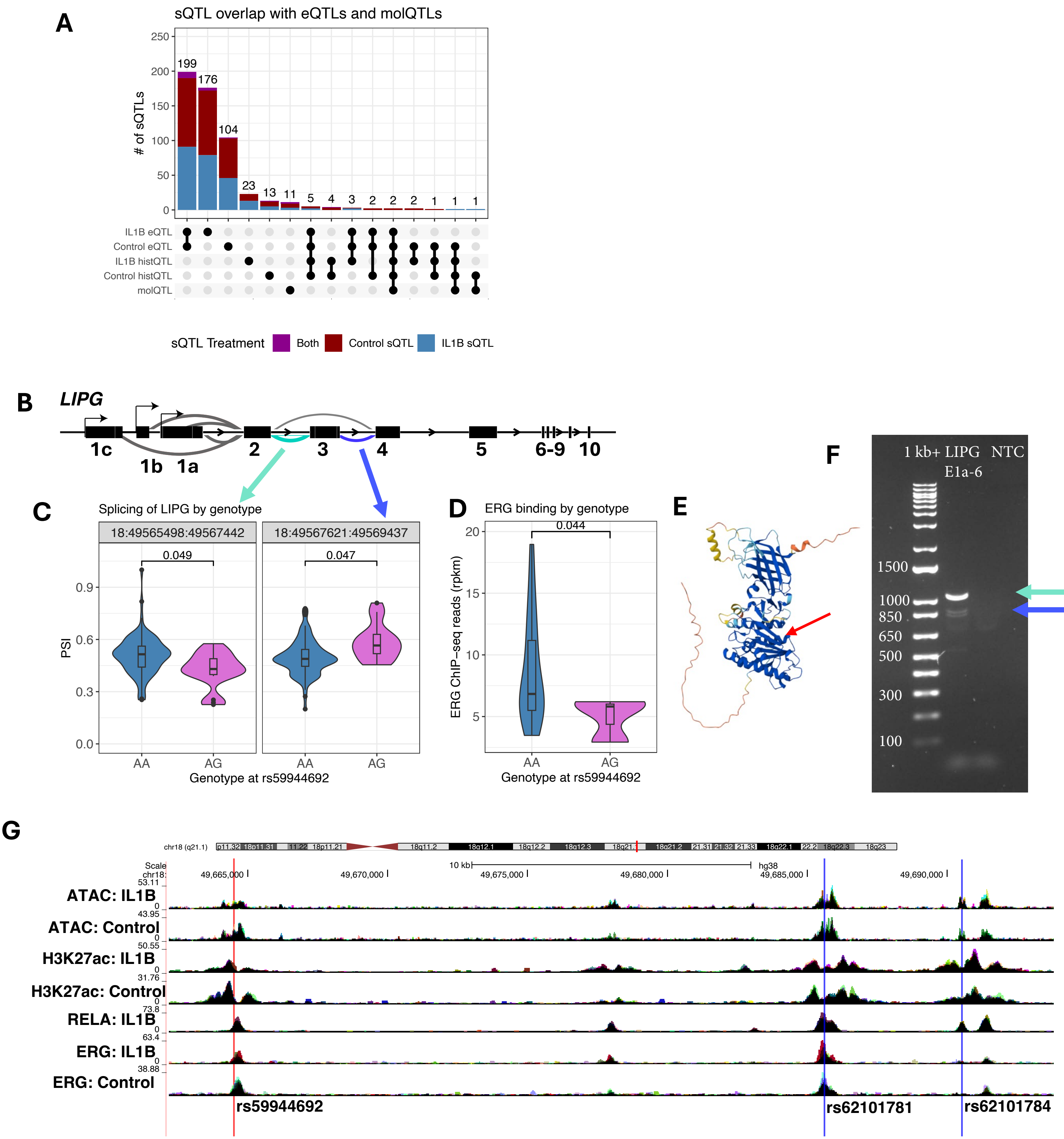


### Supplemental Figure 4

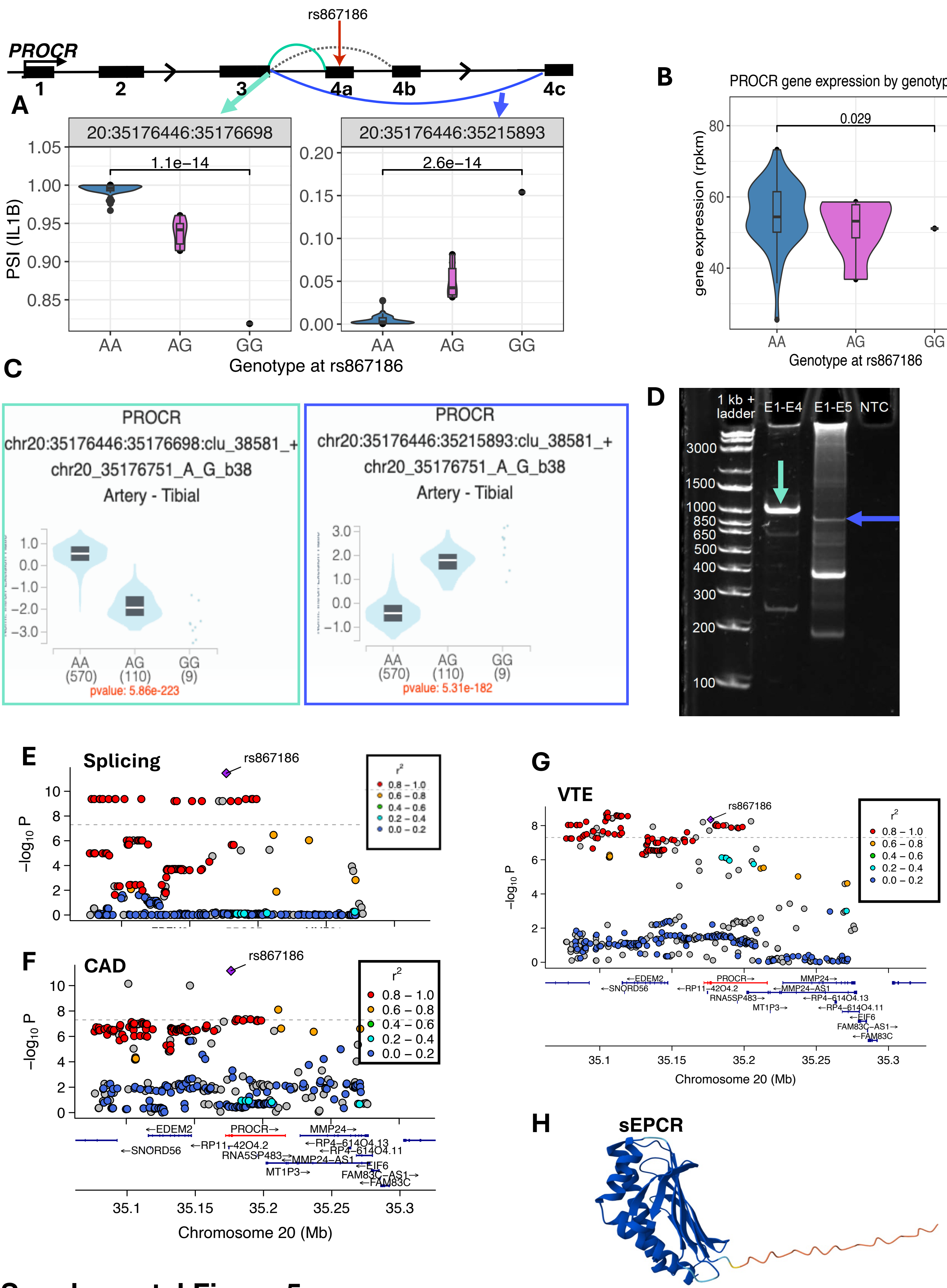


**Supplemental Figure 5**

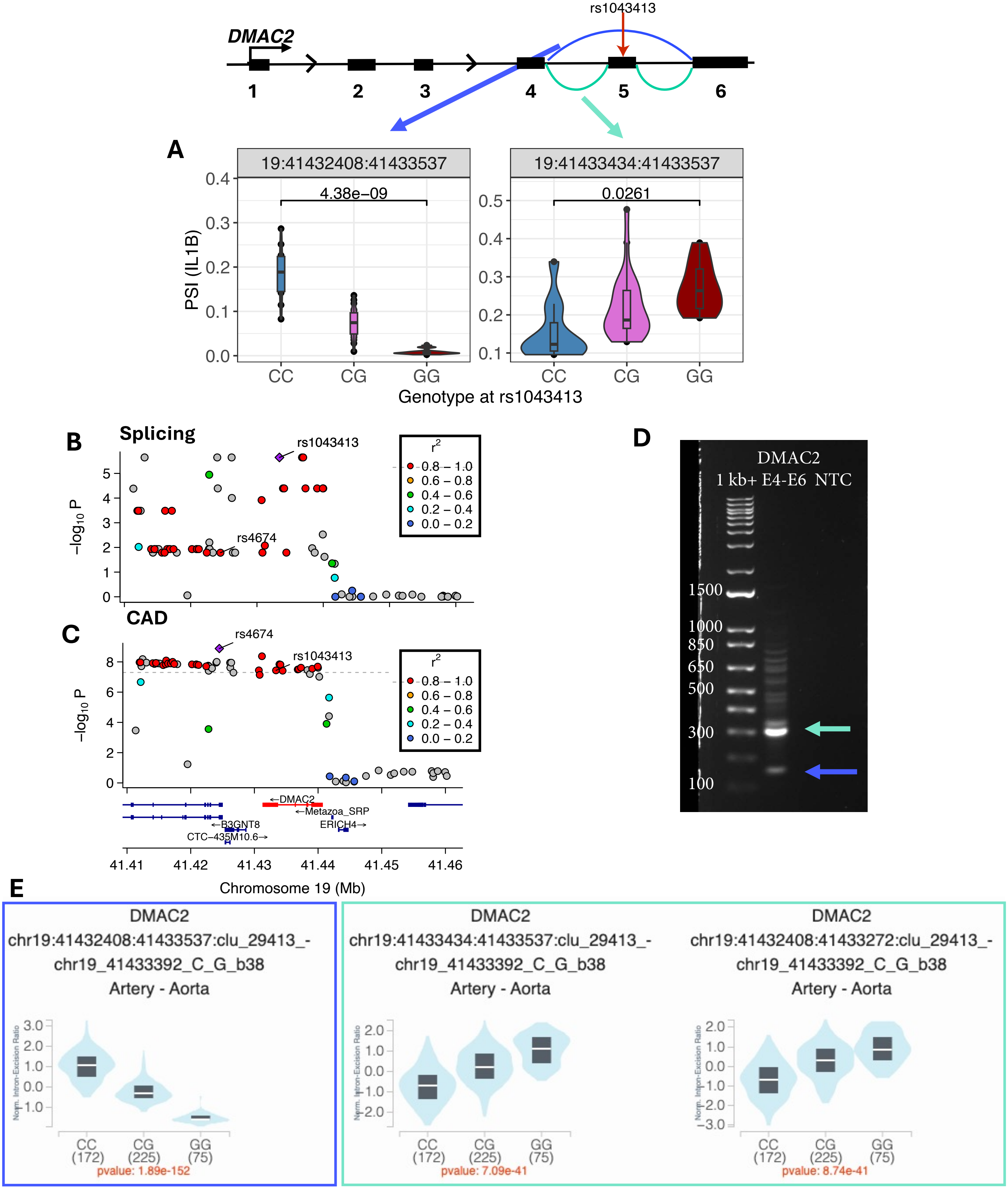


**Supplementary Figure 6**
